# ISWI1 complex proteins facilitate developmental genome editing in *Paramecium*

**DOI:** 10.1101/2023.08.09.552620

**Authors:** Aditi Singh, Lilia Häußermann, Christiane Emmerich, Emily Nischwitz, Brandon KB Seah, Falk Butter, Mariusz Nowacki, Estienne C. Swart

**Author notes:** Equal contribution (Aditi Singh, Lilia Häußermann).

## Abstract

Chromatin remodeling is required for essential cellular processes, including DNA replication, DNA repair, and transcription regulation. The ciliate germline and soma are partitioned into two distinct nuclei within the same cell. During a massive editing process that forms a somatic genome, ciliates eliminate thousands of DNA sequences from a germline genome copy in the form of internal eliminated sequences (IESs). Recently we showed that the chromatin remodeler ISWI1 is required for somatic genome development in the ciliate *Paramecium tetraurelia*. Here we describe two paralogous proteins, ICOP1 and ICOP2, essential for DNA elimination. ICOP1 and ICOP2 are highly divergent from known proteins; the only domain detected showed distant homology to the WSD motif. We show that both ICOP1 and ICOP2 interact with the chromatin remodeler ISWI1. Upon *ICOP* knockdown, changes in alternative IES excision boundaries and nucleosome densities are similar to those observed for *ISWI1* knockdown. We thus propose that a complex comprising ISWI1 and either or both ICOP1 and ICOP2 are needed for chromatin remodeling and accurate DNA elimination in *Paramecium*.

## Introduction

Chromatin’s underlying subunit, the nucleosome, is highly conserved ∼146 base pairs of DNA wrapped around a histone octamer. The presence of a nucleosome on a DNA sequence alters its geometry and physically shields DNA, affecting its interaction with other DNA-binding proteins (Morgunova and Taipale 2021; Pryciak and Varmus 1992; Piña et al. 1990). Thereby, the nucleosome regulates its participation in numerous molecular processes (Bai and Morozov 2010; Price and D’Andrea 2013; Campos and Reinberg 2009; Alabert and Groth 2012).

Nucleosomes can be moved, ejected or reconstructed with histone variants by four families of ATP-dependent chromatin remodelers (Clapier and Cairns 2009). The imitation switch (ISWI) family of chromatin remodelers forms several complexes capable of nucleosome sliding (Längst et al. 1999) in different organisms, each serving a distinct role. ISWI contains an N-terminal SNF2 ATPase domain that provides energy to move the nucleosome (Li et al. 2019). The HAND-SANT-SLIDE domain (HSS) in the C-terminus is essential for substrate recognition (Grüne et al. 2003). ISWI complex partners determine the context of the complex activity and alter its remodeling efficiency (Längst et al. 1999; Toto et al. 2014). ISWI complexes have been shown to regulate DNA replication, transcription, DNA repair, and V(D)J cleavage of polynucleosomal DNA (Clapier and Cairns 2009; Aydin et al. 2014; Patenge et al. 2004).

We recently showed that an ISWI homolog, ISWI1, is required for genome editing in *Paramecium tetraurelia* (henceforth, *Paramecium*) (Singh et al. 2022). Like other ciliates, *Paramecium* has distinct nuclei: the germline micronucleus (MICs) and the somatic macronucleus (MAC). The MICs produce gametic nuclei that form a diploid zygotic nucleus, which generates new MICs and MACs. The zygotic genome developing into a new MAC genome undergoes massive editing, excising thousands of germline-limited sequences and also genome amplification to a high polyploidy (∼800n) (Zangarelli et al. 2022; Drews et al. 2022a). *Paramecium*’s internal eliminated sequences (IESs) are distributed throughout intergenic and coding regions in the germline genome (Arnaiz et al. 2012). IESs removal requires precise excision and subsequent DNA repair, ensuring a functional somatic genome (Dubois et al. 2012; Kapusta et al. 2011).

Each of *Paramecium*’s 45,000 unique IESs is flanked by conserved 5’-TA-3’ dinucleotides, which are part of a less well-conserved ∼5 bp terminal inverted repeat (Arnaiz et al. 2012; Bischerour et al. 2018; Klobutcher and Herrick 1995). PiggyMAC (PGM), a domesticated PiggyBac transposase (Baudry et al. 2009; Bischerour et al. 2018), is responsible for the excision of IESs and other germline-specific sequences in *Paramecium*. The IES length distribution monotonically declines with a characteristic 10/11 bp periodicity, except for ∼34-44 bp “forbidden” peak, where IESs appear largely absent (Arnaiz et al. 2012). The interruption is supposedly caused by the requirement of DNA looping for the excision of longer IESs (Arnaiz et al. 2012).

Since IESs lack a well-conserved motif, additional molecules are required for their recognition and excision in addition to PGM. Germline-limited sequences are thought to be targeted by two small non-coding RNA classes: scnRNAs and iesRNAs. scnRNAs are produced by Dicer-like proteins Dcl2 and Dcl3 in the MICs and on Piwi proteins Ptiwi01/09, facilitating nuclear crosstalk and DNA elimination in the new MAC (Bouhouche et al. 2011); (Lepère et al. 2009; Sandoval et al. 2014). iesRNAs, produced by Dcl5 and Ptiwi10/11 proteins, supposedly form a positive feedback loop after the initial onset of IES excision to efficiently excise all IES copies (Sandoval et al. 2014; Furrer et al. 2017). As a general trend, shorter IESs tend to be older and primarily independent of iesRNAs and scnRNAs, whereas younger, longer IESs require additional molecules for excision (Sellis et al. 2021). In addition, Ptiwi01/09 was also proposed to interact with the PRC2 complex (Miró-Pina et al. 2022; Wang et al. 2022), repressing transposable elements, and with ISWI1, required for the IES’s precise excision (Singh et al. 2022).

The depletion of ISWI1 is lethal and leads to two distinct errors: failure of excision of numerous IESs and alternative IES excision at the wrong TA boundaries (Singh et al. 2022). In the latter case, excision precision was proposed to be compromised by inappropriate nucleosome positioning. A distinctive characteristic of ISWI1-depletion is the substantial fraction of alternatively excised IESs of the “forbidden” peak length. In this study, we identified and investigated the subunits of the ISWI1 complex and their contribution to IES excision.

## Results

### Identifying putative components of the ISWI1 complex

Previously, we performed co-immunoprecipitation (co-IP) of proteins associated with 3XFLAG-HA-tagged ISWI1 (Singh et al. 2022). After ISWI1, the most abundant protein candidate detected by mass spectrometry (MS), with more than five-fold enrichment in peptides identified relative to the input, is a 779 amino acid-long uncharacterized protein (ParameciumDB identifier: PTET.51.1.P0440186). The ohnolog of the candidate protein from its whole genome duplication (PTET.51.1.P0180124, 783 amino-acid long) is also present in the subset of peptides identified as unique to ISWI1-IP replicates in the same MS dataset (Singh et al. 2022). We characterized these proteins further to determine whether they are part of the ISWI1 core complex functioning in genome editing.

To begin, we checked if the candidate proteins have homologs that form ISWI complexes in other organisms (Dirscherl and Krebs 2004). Since Pfam database searches failed to identify any domain (Finn et al. 2003), we searched for more distantly associated domains using HHpred (Zimmermann et al. 2018). HHpred generates an HMM for the query using the iterative search and alignment functionality provided by HHblits (Remmert et al. 2011). The HHpred results indicated a probability of 91.68% for the “D-TOX E motif, Williams-Beuren syndrome DDT (WSD) motif” (Pfam model PF15613; 65 aa), located centrally in the candidates (Fig 1A & B). This motif is also present in the WHIM2 domain (InterPro ID: IPR028941), which is known to interact with linker DNA and the SLIDE domain in ISWI proteins (Aravind and Iyer 2012; Yamada et al. 2011; Mukherjee et al. 2009). Based on this analysis and subsequent experimental complex determination investigations, we named our putative interacting candidates ISWI1 Complex Protein 1 (ICOP1) and its closely-related ohnolog ISWI1 Complex Protein 2 (ICOP2).

**Figure 1:**
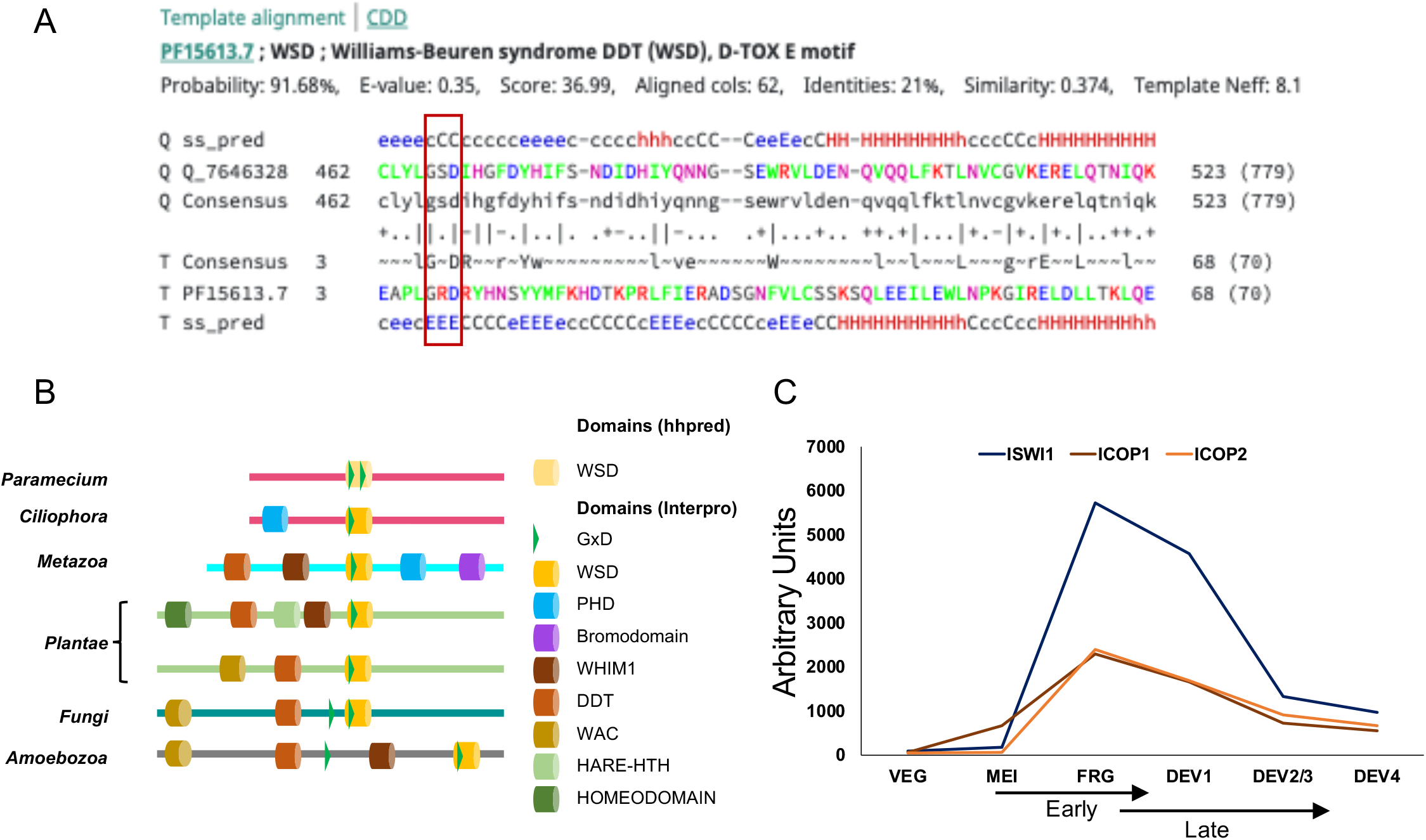
Identification of ISWI Complex Proteins (ICOP). (A) Template alignment generated by HHpred analysis of ICOP1 showing 91.68% probability match (E-value 0.35) with Williams-Beuren syndrome DDT(WSD) or D-TOX E motif. The conserved GxD signature is highlighted with a red bar.; Q= Query (ICOP1); ss_pred: secondary structure prediction; T= template (B) Representative domain architecture of WHIM2 domain-containing proteins used to create phylogeny. (C) mRNA expression profile (arbitrary units) of ICOP1 and ICOP2 in comparison to ISWI1 during autogamy. VEG: vegetative, MEI: the stage where MICs undergo meiosis and maternal MAC begins to fragment, FRG: about 50% of cells with fragmented maternal MAC, Dev1: the earliest stage with visible developing macronuclei (anlage), Dev2/3: most cells with macronuclear anlage, Dev4: most cells with distinct anlage. MEI and FRG constitute the “Early” time point, and the “Late” time point consists of Dev1 and Dev2/3 stages.

*ICOP1* and *ICOP2* are upregulated during autogamy and have an expression profile similar to *ISWI1*’s (Fig 1C). Generally, proteins with WHIM2 domains have multiple domain architectures (Aravind and Iyer 2012). ICOP1 and ICOP2 proteins had no additional conserved domains except for the three amino acid residues, called the GxD signature (Figs 1A & B), within the identified WSD motif. Furthermore, our phylogenetic analysis of proteins with the WSD motif suggests that ICOP1 and ICOP2 are highly divergent in comparison to other WSD motif-containing proteins (Ext. Fig 1).

### ICOP proteins localize to the developing MACs during autogamy

Since ISWI1-GFP localizes in the developing MAC during autogamy (Singh et al. 2022), we examined the localization of the ICOP proteins. We co-transformed paramecia with either N-terminally tagged HA-ICOP1 or C-terminally tagged ICOP2-HA with ISWI1-GFP and observed that all these proteins localized exclusively to the developing MACs during autogamy (Fig 2A). We observed no growth defects in the co-transformed cells during vegetative growth or in the F1 progenies (Ext. Fig 2A). Their localization suggests that ICOP paralogs and ISWI1 function at the same stages during new MAC development.

**Figure 2:**
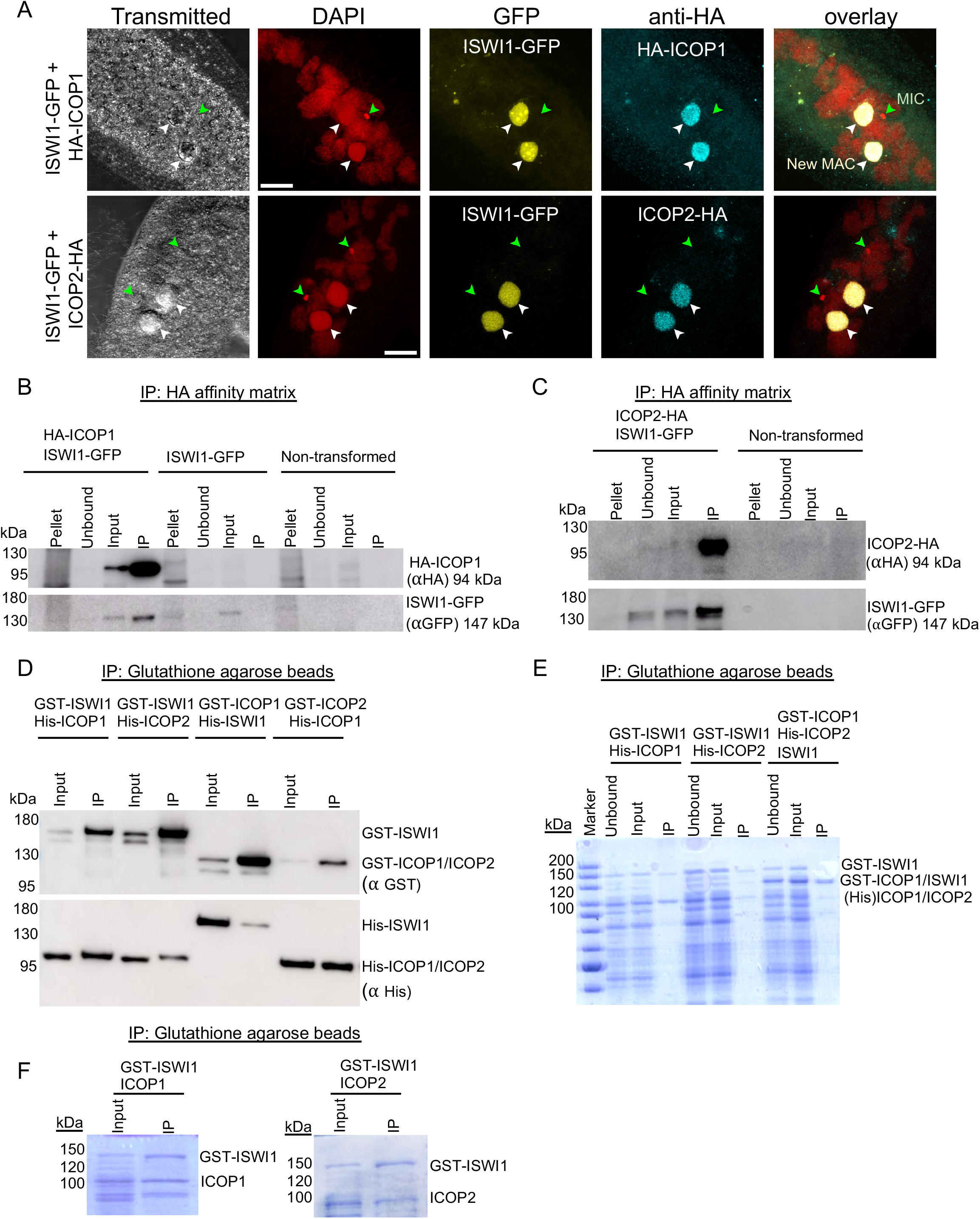
Interaction of ICOP1 and ICOP2 with ISWI1 in new MACs. (A) Confocal fluorescence microscopy images of HA-ICOP1, ICOP2-HA, and ISWI1-GFP localization: maximum intensity projections of z-planes. Red =DAPI. Yellow =GFP. Cyan =HA. Green arrow =MIC. White arrow = new MAC. All channels were optimized individually for the best visual representation. DAPI channel of ICOP2-HA: Gamma factor = 0.8. Scale bar = 10 µm. (B) & (C) Western blot, co-immunoprecipitation (co-IP) of HA-ICOP1/ISWI1-GFP and ICOP2-HA/ISWI1-GFP in *Paramecium*. Controls: non-transformed and ISWI1-GFP transformed. (D-F) co-IP after *E. coli* expression and pulldown; (D) Western blot, (E&F) Coomassie staining. GST-ISWI:147 kDa, His-ISWI1:122 kDa, His-ICOP1/2:95 kDa, GST-ICOP1/ICOP2:119 kDa, untagged ISWI1:120 kDa, untagged ICOP1/2:93 kDa.

### ISWI1 and ICOP paralogs form a complex *in vivo* during autogamy

Using the co-transformed HA-ICOP1/ISWI1-GFP or ICOP2-HA/ISWI1-GFP lysates, we performed reciprocal co-IPs to assess ICOP1 and ICOP2 interactions with ISWI1. As controls, wild-type, non-transformed, and only ISWI1-GFP transformed lysates were used. As expected, non-transformed cells showed no protein pulldown signal with either HA- or GFP-conjugated beads (Figs 2B & 2C, Ext. Fig 2B). ISWI1-GFP signal was detected only in the “input” fraction when using the HA-conjugated beads (Fig 2B, lower panel) in the single transformants. ISWI1-GFP was successfully co-purified with HA-ICOP1 or ICOP2-HA from the co-transformed cell lysates (Figs 2B & C, and Ext. Fig 2B). co-IPs with ISWI1-GFP, HA-ICOP1, and ICOP2-HA single transformants were analyzed using MS (Ext. Fig 2B & C). ISWI1 was among the most highly enriched proteins, along with either one or both of the ICOPs in MS (Ext. Fig 2D). Therefore, we conclude that both ICOP paralogs can interact with ISWI1 in *Paramecium*.

### ICOPs do not require a GxD signature for interaction with ISWI1

Since ICOP1 and ICOP2 are part of the ISWI1 complex, we investigated whether the paralogs can bind directly to ISWI1 by co-expressing ICOP1, ICOP2, and ISWI1 in *E. coli*. N-terminal fusion (GST or His) or untagged proteins were used for the pulldown. First, we validated the specificity of the pulldowns using either glutathione agarose (GST) beads or nickel-IMAC agarose (Ni_2_+NTA) beads. We did not observe unspecific binding or cross-reactivity of tagged proteins in the IP fraction of the pulldowns (Ext Fig 2E-G). Next, we co-expressed ISWI1, ICOP1, and ICOP2 in different combinations and performed pulldowns using GST beads. The three proteins were pulled down together, suggesting they have a direct affinity for each other (Fig 2D-F).

Since the GxD signature in WHIM-containing proteins was proposed to mediate interactions with ISWI1 in diverse eukaryotic organisms (Aravind and Iyer 2012), we assessed whether this signature is needed to form the ISWI1-ICOP complex. ICOP1 and ICOP2 have two GSDs (Fig 3A); however, only the first one aligns with the HMM GxD (Fig 1A). Aspartate was proposed as the essential driver of the interaction in the GxD signature (Aravind and Iyer 2012). We generated ICOP mutants with either a D to A substitution (GxA mutants) or the complete deletion of GxD (delGSD mutants) (Fig 3B). His-ISWI1 co-purified with GST-ICOP mutant proteins, albeit somewhat less than the wild-type proteins (Fig 3C). Nevertheless, our data indicates that the ISWI1 and ICOPs could interact without the GxD signature.

**Figure 3:**
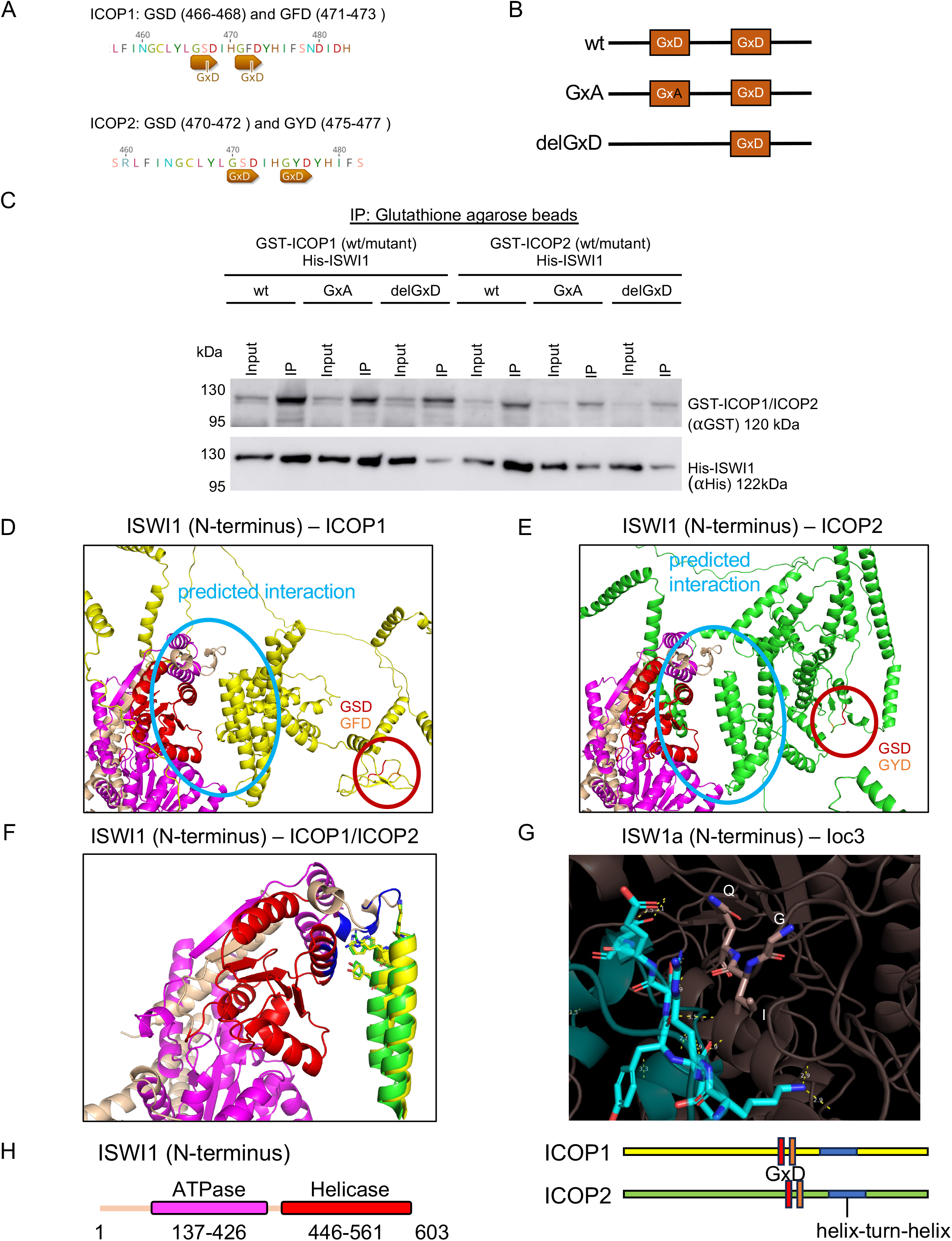
Investigation of the GxD signature in ICOP/ISWI1 interaction. (A) Screenshots from Geneious Prime (version 2023.1.1) showing GxDs in ICOP1 and ICOP2, annotated in brown. (B) Schematic representation of GxD mutants generated in this study. (C) Western Blot on co-IP of GST-ICOP GxD mutants and His-ISWI1 overexpressed in *E*. *coli* probed with by anti-GST and anti-His antibodies; GST-ICOP wild-type is used as control. (D-F) Structure prediction of multimers (ISWI1 N-terminus (residues 1-603) with ICOP1 or ICOP2) with AlphaFold (version 2.2.0). ICOP1: yellow, ICOP2: green, GSD signature: red, GFD/GYD: orange, ISWI1: wheat, ISWI1 ATPase domain: magenta, ISWI1 helicase domain: red. (D) & (E) ISWI1-ICOP1 and ISWI1-ICOP2 interaction, respectively. Predicted interaction interface is highlighted with blue circles. Both GxDs are highlighted with red circles. (F) ISWI1 N-terminus with interacting helices of ICOP paralogs (ICOP1: residues 556-597; ICOP2: residues 560-603). Proximate residues on ISWI1 are shown in blue. Proximate residues of ICOPs are shown as sticks. (G) GxD signature in the published crystal structure (PDB accession number 2Y9Y): ISW1a (del_ATPase; cyan) and Ioc3 (WHIM containing protein; dark salmon) from yeast. GxD signature (GIQ in Ioc3) and spatially close residues in ISW1a are shown as sticks, polar contacts in yellow. (H) Schematic representation of the sequences used for predictions in (D) & (E).

We predicted the interaction of ISWI1 and ICOPs using AlphaFold2. ISWI1’s predicted structure was of high confidence, and its domains showed similarity to published structures from yeast (Ext. Fig 3A & B). ICOP structures had low confidence, most likely due to their high divergence from other known structures (Ext. Fig 3B). For the complex prediction, AlphaFold2 version 2.3.0 predicted interactions in all tested combinations with large interaction interfaces, while version 2.2.0 predicted an interaction of either ICOP1 or ICOP2 only with the N-terminus of ISWI1 (residues 1-603, including the ATPase domain but not the HSS domain) (Fig 3D-F). In these models, the ICOPs bound with a defined helix-loop-helix motif (ICOP1: residues 556-597; ICOP2: residues 560-603) (Fig 3F). Irrespective of the AlphaFold2 version, neither of the GxD signatures were predicted to participate in the interaction (Fig 2D & E, Ext. Table 1).

### *ICOP1/2*-KD affects cell survival and genome editing

*ICOP1* and *ICOP2* were knocked down by RNAi, either individually or together, to assess their role in genome editing. Knockdown of *ND7*, a gene involved in trichocyst discharge (exocytosis) (Skouri and Cohen 1997), was used as negative control (CTRL). Previously published *ISWI1*-KD data (Singh et al. 2022) was used as positive control and for comparative purposes. The efficiency of the different knockdowns (KDs) was confirmed using RNA-seq: in all KD cases, the expression of the target gene was substantially reduced compared to the controls (Fig 4A). Allowing no mismatches, the off-target tool on ParameciumDB predicted a 24 bp window in *ICOP2* that can be co-silenced with the *ICOP1* RNAi construct (*Paramecium* siRNAs are typically 23 nt). *ICOP1* mRNA levels were reduced in *ICOP2*-KD and vice versa, but not to the extent of the RNAi targets (Fig 4A). *ICOP1*-KD led to 30% lethality, while *ICOP2*-KD led to about 20% lethality, and a double KD of *ICOP1* and *ICOP2* led to about 65% lethality in the F1 generation (Fig 4B). Additionally, most cells in the single knockdowns failed to grow at a standard division rate (“sick” cells; Fig 4B).

**Figure 4:**
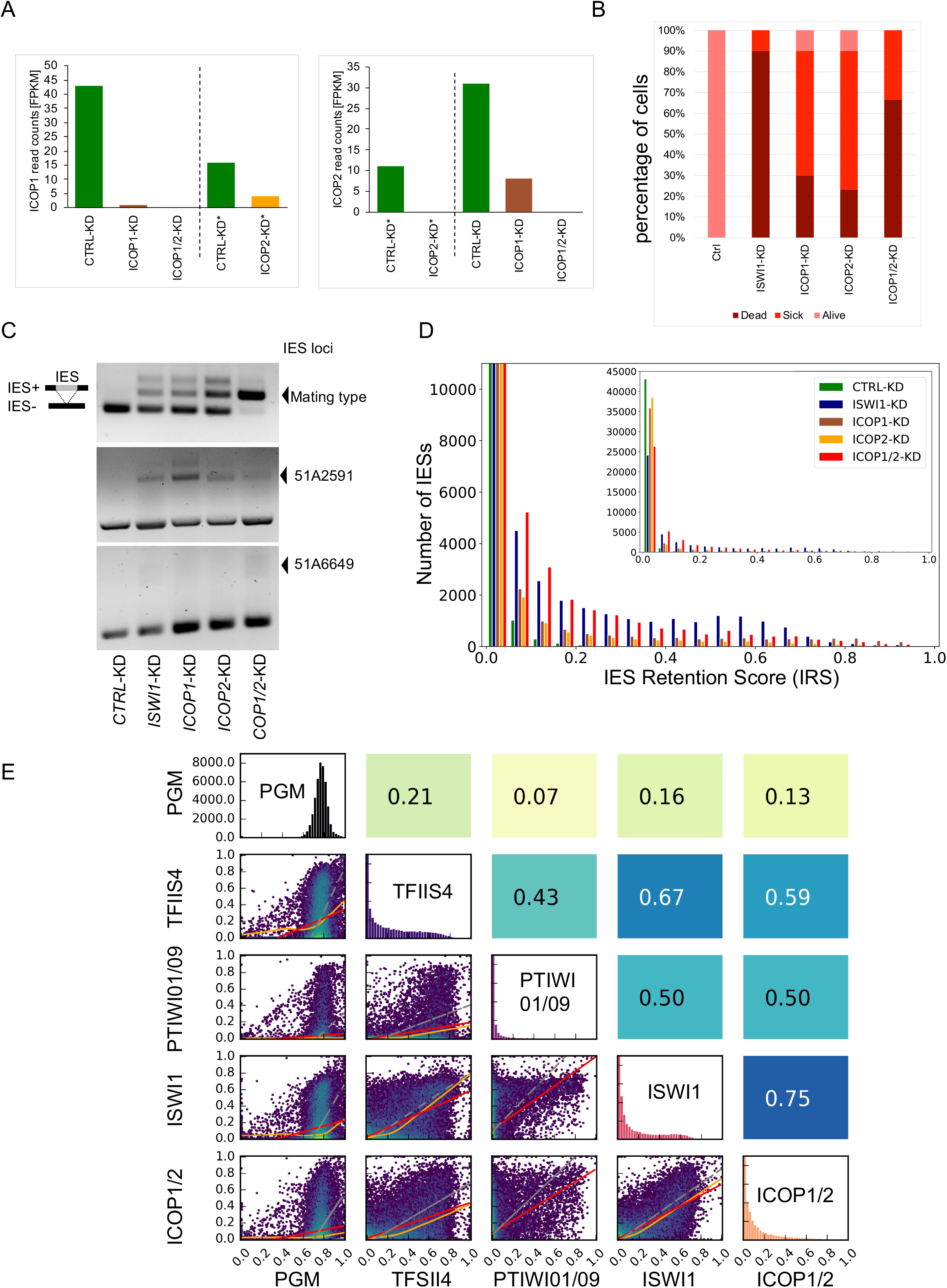
Effects of *ICOP* knockdowns on DNA excision. (A) mRNA expression levels in FPKM (Fragments per kilobase per million mapped reads) compared between knockdowns for ICOP1 and ICOP2 transcripts early in development (40% old MAC fragmentation) or asynchronous culture (*). (B) Survival of recovered post-autogamous knockdown cells followed for several vegetative divisions. Alive (pink): normal division. Sick (red): slower division rate. Dead (cayenne): no cells. (C) Retention of individual IESs, *ISWI1*-KD = positive control. Retained IESs (IES+) result in a larger amplicon. (D) Genome-wide IES retention in different KDs. Histogram of IES retention scores (IRS = IES+ reads/(IES+ reads + IES-reads)). (E) Correlation of IRSs among KDs. Diagonal: IRS distributions of individual KDs. Below diagonal: correlation graphs of pairwise comparisons. Above diagonal: corresponding Pearson correlation coefficients. Red lines: ordinary least-squares (OLS) regression, orange lines: LOWESS, and gray lines: orthogonal distance regression (ODR).

With PCRs on known IES loci, we checked whether the *ICOP* KDs affect IES excision (Fig 4C). Longer fragments containing IESs (IES+) were amplified in all KD permutations, suggesting ICOPs are essential during genome editing. Next, we investigated the genome-wide effect of *ICOP* KDs. The IES retention score (IRS) was calculated for each IES to study the global effect on IES excision. Both single and double KDs caused IES retention, with a stronger effect in *ICOP1/2*-KD (Fig 4D). Like *ISWI1*-KD, *ICOP1/2*-KD IRSs correlated modestly with IRSs of other gene KDs known to affect IES excision (e.g., Fig 4E).

### *ICOP1/2*-KD affects IES excision precision

Errors in IES excision manifest not only as IES retention but also as imprecise IES excision. Imprecise or alternative excision in *Paramecium* occurs naturally at TA dinucleotides that are not the predominant IES boundaries (Duret et al. 2008) (Fig 5A). Generally, alternative excision occurs at low levels in nature (CTRL-KD, Fig 5B & C). *ISWI1*-KD substantially enhances alternative excision versus KDs of other genome-editing genes (Singh et al. 2022). Similar to *ISWI1*-KD, *ICOP1*-KD and *ICOP2*-KD elevate imprecise excision, though to a lesser extent in both single and double KDs (Fig 5B, Ext. Table 2). Previously (Singh et al. 2022), we did not measure the IESs where 100% of the mapped reads were alternatively excised (Ext. Table 2), thus underestimating alternative excision. Nevertheless, by the old estimation method, the percentage of alternative excision events per IES was highest in *ICOP1-*KD (mean 7%) and similar between *ICOP2*-KD (mean 4.2%) and *ICOP1/2*-KD (mean 4.7%). This is higher compared to the other KDs (mean range 1.5-2.4% (Singh et al. 2022)) except *ISWI1*-KD (mean 9.2% (Singh et al. 2022); Ext. Table 2).

**Figure 5:**
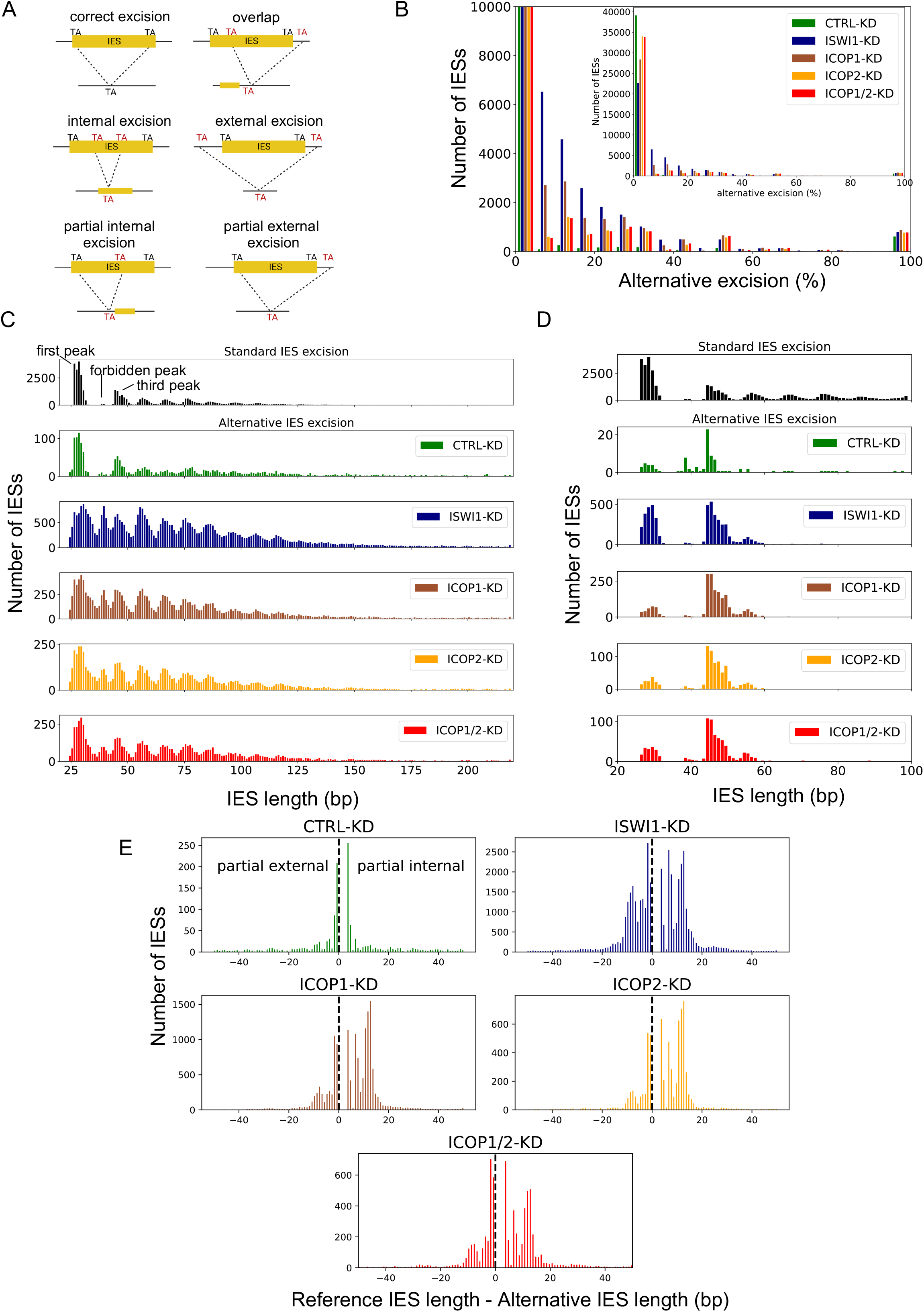
Alternative IES excision in *ICOP* knockdowns. Analysis for *ISWI1*-KD, *ICOP1*-KD, *ICOP2*-KD, and *ICOP1/2*-KD, with *ND7*-KD as the negative control. (A) Schematic representation of analyzed IES excision events. (B) Distribution of genome-wide alternative IES excision (percent per IES) for different KDs. (C) Length distribution of alternatively excised IESs for each KD. The reference length distribution for all IESs is given above (“Standard IES excision”). (D) Origin of alternatively excised IESs in the “forbidden” peak. The reference length is plotted for all alternatively excised 34 – 44 bp IESs. (E) Length distribution of partial external and partial internal alternative excision events for the KDs.

The use of alternative TA boundaries changes the length of the excised fragments. The maximum and minimum length of excised IESs was shifted towards more extremes, and generally, alternatively excised IESs were longer than the reference length (Ext. Table 3). The length distribution of alternatively excised IESs resembled the ∼10 bp periodicity characteristic of *Paramecium* IESs, with the striking exception that the “forbidden” peak (Arnaiz et al. 2012) was present in all three *ICOP* KDs, as in *ISWI1*-KD (Fig 5C). In *ISWI1*-KD, alternative IESs in the “forbidden” peak mainly originated from the first and third peaks, while they primarily originated from the third peak in *ICOP* KDs (Fig 5D). The similarity in alternative excision effects of *ISWI1* and *ICOP* KDs suggests that ISWI1 and ICOP proteins cooperate in the precise excision of IESs.

Further, we examined five possible alternative IES excision events: “partial internal”, “partial external”, “overlap”, “internal,” and “external” (Fig 5A). Generally, “internal” and “external” are low-frequency events in all KDs (Ext. Fig 4A). In control KD, “overlap”, “partial external” and “partial internal” events were approximately equal at around 30% each (Ext. Fig 4B). This contrasts with *ICOP*s and *ISWI1* KDs, where “overlap” was relatively infrequent, while “partial internal” and “partial external” comprised the largest share of erroneous excision events (Fig 5E, Ext. Fig 4B, Ext. Table 4). In *ISWI1*-KD, “partial internal” (-43%) and “partial external” (42%) events contributed equally, while “partial internal” dominated the *ICOP* KDs. The preference was more pronounced in the single KDs (“partial internal” - 57%; “partial external” - 28% for *ICOP1-* and *ICOP2-*KD) than in *ICOP1/2*-KD (“partial internal” - 47%; “partial external” - 34%) (Ext. Fig 4B).

### *ICOP1/2*-KD does not alter ISWI1 localization but affects scnRNAs and iesRNAs

We knocked down *ICOP1* and/or *ICOP2* to check whether their expression is required for the localization of ISWI1-GFP. As in control cells with no RNAi (Fig 6A), ISWI1-GFP localization was not impaired in *ICOP* KDs (Fig 6C-E). Only in *ISWI1*-KD, the GFP signal was entirely lost from the new MAC (Fig 6B). In *Paramecium*, the excision of a subset of IESs is suggested to depend on scnRNAs (Garnier et al. 2004). We tested the dependence of ISWI1-GFP localization on genome scanning by knocking down *PTIWI01/09*, a core protein of the scanning pathway. ISWI1-GFP localized to the new MAC upon *PTIWI01/09*-KD (Fig 6F). This suggests ISWI1 localization is independent of ICOP(s) and genome scanning.

**Figure 6:**
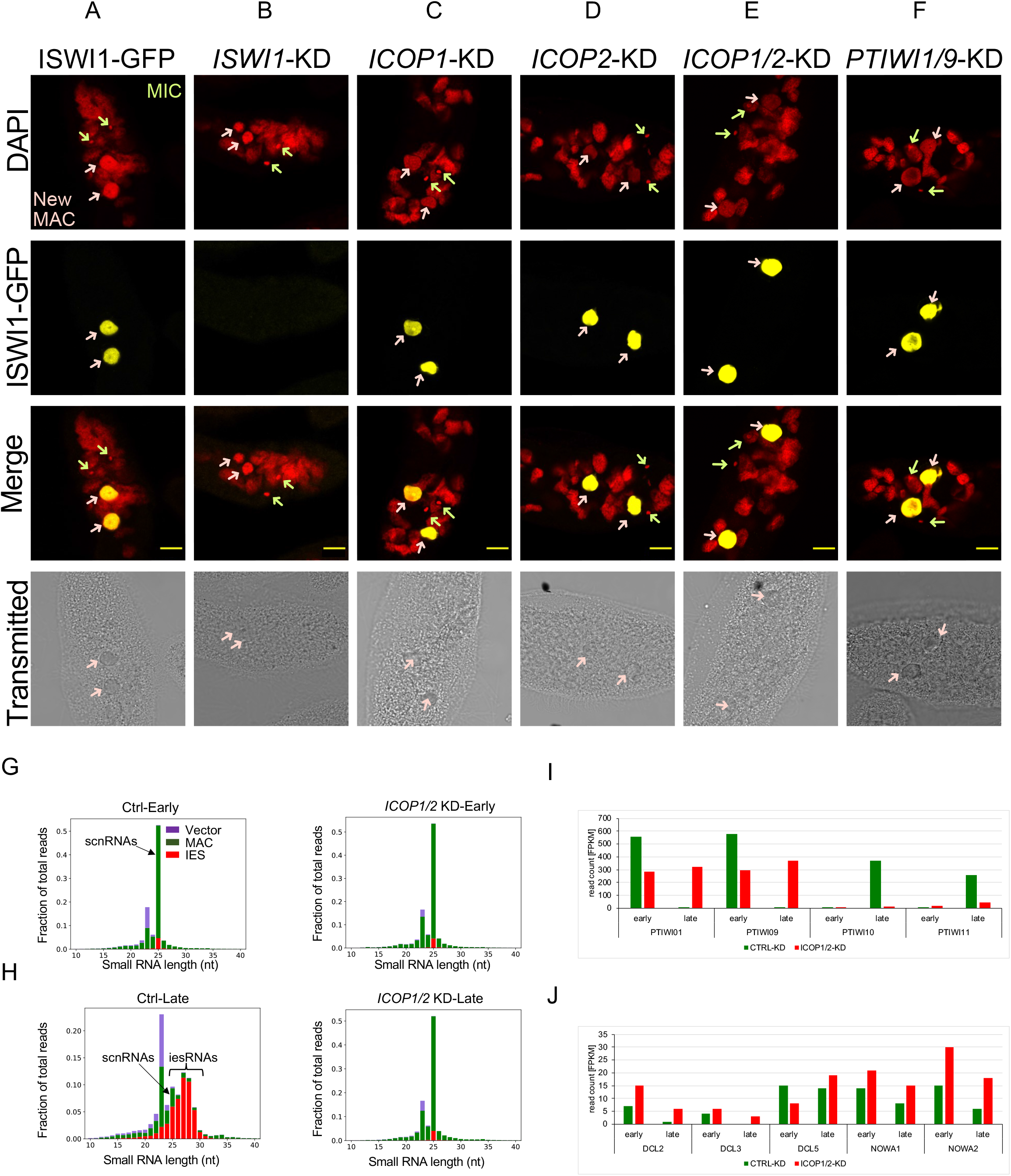
Effects of ICOP1 and ICOP2 knockdowns on ISWI-GFP localization, sRNAs and gene expression. (A-F) Confocal fluorescence microscopy of ISWI1-GFP localization under gene knockdowns. (A) Positive control: ISWI1-GFP transformed cells without RNAi; Red = DAPI, Yellow = GFP. Green arrow = MIC; pink arrow = new MAC, scale bar = 10 µm. (G & H) Histogram of 10 to 40 nt sRNAs. sRNA reads were mapped to the L4440 plasmid sequence (Vector, purple), macronuclear genome (MAC, green), and IESs (IES, red). Early = 40% of cells have fragmented MAC, Late = most cells with visible new MAC. (I & J) Histogram of mRNA expression levels in FPKM (Fragments per kilobase per million mapped reads) for different developmental-specific genes.

Next, we checked whether *ICOP1/2*-KD influences the small RNA population. scnRNAs are generated in MICs well before the development of new MACs (Lepère et al. 2009). Consequently, their production is only affected by genes involved in their biogenesis. As expected, in early development, we did not observe a pronounced effect on scnRNA production in *ICOP1/2*-KD compared to the control *ND7*-KD (*Ctrl*-KD) (Fig 6G). Knockdowns of genes whose proteins localize and function in genome editing inhibit iesRNA production by blocking the positive feedback loop for further IES excision (Allen et al. 2017). We observed the same for *ICOP1/2*-KD (Fig 6H).

Comparing the MAC-matching scnRNAs relative to the siRNAs, it is clear that there was a greater quantity of MAC-matching scnRNAs in the late time point for *ICOP1/2*-KD than for *Ctrl*-KD. This suggests that the removal of MAC-matching scnRNAs, as proposed by the RNA scanning model, was impaired by *ICOP1/2*-KD (Fig 6H). We examined sRNA biogenesis-related gene transcription in *ICOP1/2*-KD vs the control KD (Fig 6I & J). In the late developmental stages, when the ICOPs localize to the new MAC, *PTIWI10* and *PTIWI11* expression was almost completely lost upon *ICOP1/2*-KD (Fig 6I); expression of *PTIWI01, PTIWI09*, *DCL2*, *DCL3* and *NOWA1/2* was upregulated (Fig 6J).

### *ICOP1/2*-KD IES nucleosome density changes are similar to those of *ISWI1*-KD

To further investigate the functional contribution of the *ICOP* paralogs to the ISWI1 complex, we analyzed the effects of *ICOP* KDs on IES nucleosome densities. IESs with high retention in *ICOP1/2*!”# %&’( ) *+,- ./01/1 .2 345/ 3673/8 09:;/2<2=/ 1/0<6.6/< %Fig 7A) in both *ICOP1/2/PGM*-KD and *CTRL/PGM*-KD, similar to our previous observations with other knockdowns (Singh et al. 2022). The nucleosome density differences (experiment-control) for *ICOP1/2/PGM*-KD and *ISWI1/PGM*-KD had similar distributions with a narrow peak centered around 0 (Fig 7B, Ext. Table 5).

**Figure 7:**
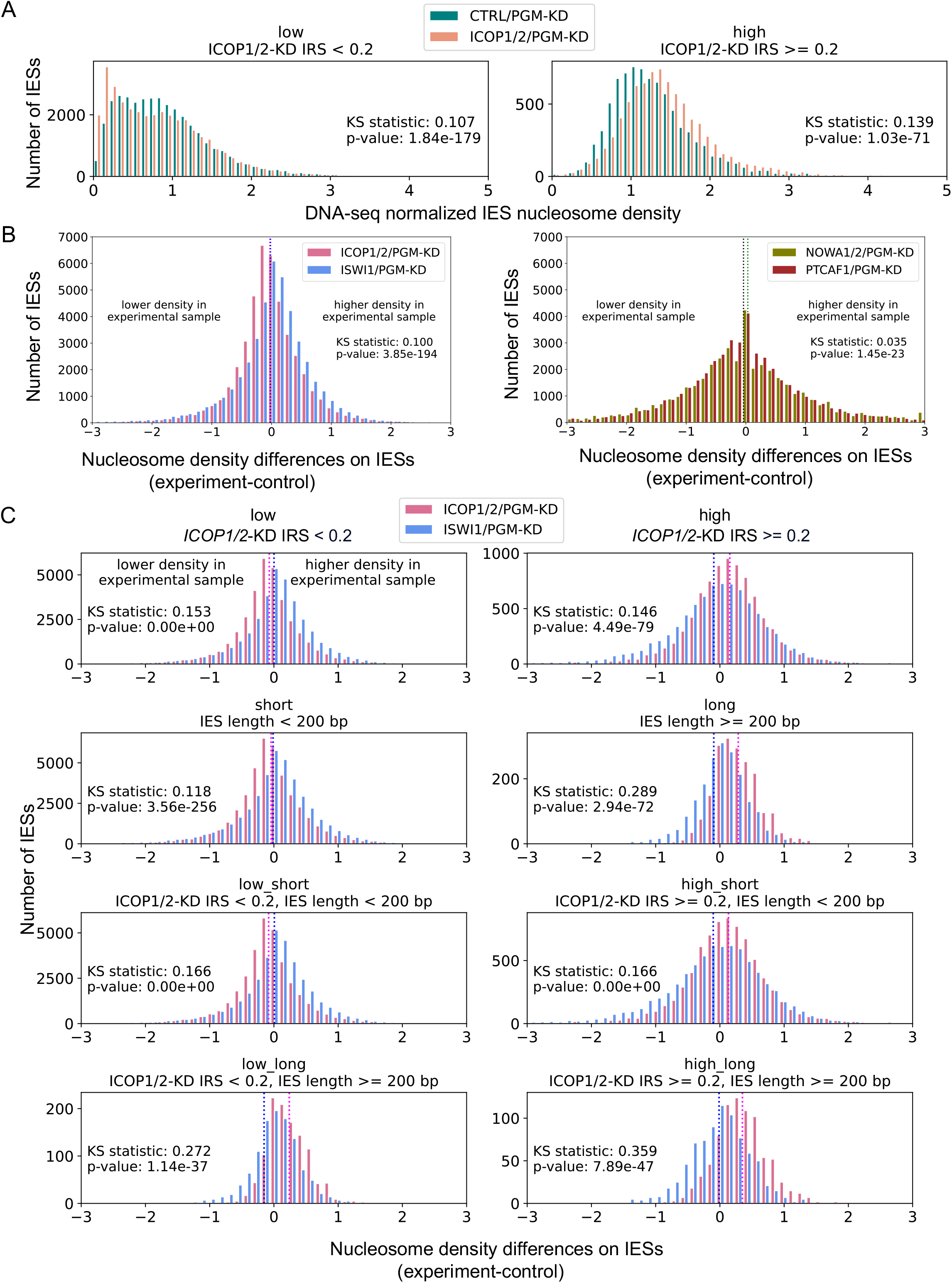
Nucleosome density changes associate with *ICOP* knockdowns. (A) Normalized nucleosome densities on IES for *ICOP1/2/PGM*-KD and *CTRL/PGM*-KD. IESs are grouped as low (IRS < 0.2) or high (IRS ≥ 0.2) according to IRSs in *ICOP1/2*-KD. (B) Nucleosome density differences for all IESs. Means are dashed lines (*ICOP1/2/PGM*-KD: magenta; *ISWI1/PGM*-KD: blue; *NOWA1/2/PGM*-KD: green; *PTCAF1/PGM*-KD: black). (C) Comparison of *ICOP1/2/PGM*-KD and *ISWI1/PGM*-KD in selected IES groups: IESs were grouped by IES retention score (IRS) in *ICOP1/2*-KD (low: IRS < 0.2; high: IRS ≥ 0.2) and IES length (short: IES length < 200 bp; long: IES length ≥ 200 bp). IES group is given above the diagrams. Means are dashed lines (*ICOP1/2/PGM*-KD: magenta; *ISWI1/PGM*-KD: blue).

However, the distributions for *NOWA1/2/PGM*-KD and *PTCAF1/PGM*-KD, which are not known chromatin remodeling proteins, were similar to each other but clearly differ from *ICOP1/2/PGM*-KD (Fig 7B). This suggests distinct effects of the remodeling complex components on nucleosome densities.

Next, IESs were grouped according to their length and IRS in *ICOP1/2*-KD. In *ICOP1/2/PGM*-KD and *ISWI1/PGM*-KD, nucleosome density differences were most prominent for long and/or ICOP1/2-dependent IESs (Fig 7C). In the *ISWI1/PGM*-KD, there was no clear trend towards higher or lower nucleosome densities, whereas, in *ICOP1/2/PGM*-KD, there tended to be higher nucleosome densities in the experimental sample (Fig 7C & Ext. Table 5). This shift towards higher nucleosome densities was also observed for *PTCAF1/PGM*-KD (Ext. Fig 6, Ext. Table 5), indicating this effect is not specific to components of the chromatin remodeling complex.

## Discussion

In this study, we identified and analyzed the role of two subunits, ICOP1 and ICOP2, that, together with the ISWI1 protein, form a complex in *Paramecium* and are required for genome editing and the development of a functional somatic genome.

ICOP1 and ICOP2 appear to be highly divergent from other proteins and did not have homology or domains that routine search methods could detect. One possible reason is that most Pfam domain model seeds comprise sequences from distant relatives of ciliates (animals, plants, and fungi). In such cases, it is helpful to use software like HHpred which uses a pairwise comparison of Hidden Markov Models (HMMs) that enables distant homology searches (Zimmermann et al. 2018). Thus, we identified a highly divergent WSD motif in ICOP1 and ICOP2 (Fig 1A & C). This motif is found in proteins that are subunits of the ISWI complex in several organisms (Toto et al. 2014).

Using overexpression in *Paramecium* and *E. coli*, we showed that ISWI1 formed a complex with the ICOP paralogs (Fig 2). The observations in *E. coli,* which lacks other *Paramecium* proteins, support direct binding between these proteins without any mediator or complex partner. Even though ISWI1 co-immunoprecipitates with both paralogs, ICOP2 was not substantially enriched in HA-ICOP1 co-IP, and ICOP1 enrichment in ICOP2-HA co-IP is also low (Ext. Fig. 2D). Thus, despite their ability to interact directly *in vitro*, it is likely that the ISWI1 might typically form complexes with either ICOP1 or ICOP2 subunits. The aspartate of the GxD signature in WSD is proposed to determine the interaction between ISWI and WHIM-containing proteins (Aravind and Iyer 2012). However, to our knowledge, no supporting experimental evidence exists for this suggestion. In the crystal structure, the GxD signature (GIQ) of Ioc3, a WHIM-containing complex protein of yeast ISW1a, lacks the acidic residue and forms no polar interactions with ISW1a (Fig 3G). Our heterologous expression studies show that mutation or deletion of the GxD signature does not completely abolish ICOP-ISWI interaction (Fig 3C). Furthermore, AlphaFold2 modeling predicted the interaction of ICOP paralogs at the N-terminus of ISWI1, mediated by a helix-turn-helix motif and not the GxD signature (Fig 3F & G). In the future, better structural prediction software and experimental structure determination approaches will be needed to determine precisely how the proteins interact in this complex.

Along with strong inhibition of iesRNAs, *PTIWI10*/*11* expression was abolished by the *ICOP* KDs. As these genes are transcribed in the developing MAC, the loss of *PTIWI10/11* expression could either be due to the retention of an IES in their promoter region or to nonsense-mediated decay (NMD) of mRNA triggered by IES retention in the CDS (Bazin-Gélis et al. 2023; Sandoval et al. 2014; Furrer et al. 2017). sRNA sequencing also revealed that the MAC-specific scnRNAs are elevated in *ICOP1/2-*KD compared to the control (Fig 6H). The same phenomenon has been observed in *NOWA1/2*-KD (Swart et al. 2017) and *PTCAF1*-KD (Ignarski et al. 2014). *NOWA1/2* is involved in genome scanning (Nowacki et al. 2005), whereas *PTCAF1* is a part of the PRC2 complex needed for H3K27me3 deposition during IES excision (Ignarski et al. 2014; Wang et al. 2022; Miró-Pina et al. 2022). Previously, elevated levels of MAC-specific scnRNAs were suggested as being due to inhibition of their elimination (Ignarski et al. 2014). With the caveat of the lack of replicates, we observed that, unlike *PTIWI10/11,* genes associated with scnRNAs, notably *PTIWI01*/*09*, are modestly upregulated in the late developmental stage upon *ICOP1/2*-KD, likely inhibiting MAC-matching scnRNAs from degradation. In the future, it would be worth investigating the expression of *PTIWI01*/*09* and related genome editing genes (e.g., *NOWA1/2* and *PTCAF1*) for knockdowns to observe if their expression changes are similar to those in *ICOP1/2*-KD. However, it is clear that the IES retention in *ICOP1/2*-KD is substantially stronger than the *PTIWIs* (Fig 4) and also exhibits enhanced alternative excision properties (Fig 5). Thus, altered expression levels of the *PTIWIs* and other genome editing genes cannot account for most of the observed effects in *ICOP1/2*-KD, irrespective of whether the development-specific sRNA levels or their MAC:IES ratios are altered.

Most IESs are likely remnants of autonomous or non-autonomous transposons (Seah et al. 2023; Sellis et al. 2021) that decayed beyond recognition with time due to a lack of selection pressure caused by their efficient removal during MAC genome development (Sellis et al. 2021). A third of all IESs are 26 to 28 bp in length and are proposed to be short enough to allow the interaction of two PGMs without DNA bending (Arnaiz et al. 2012). Longer IESs require DNA looping, causing 34 to 44 bp IESs in the “forbidden” peak to be highly underrepresented, either too long for two PGM subunits to interact or too short for DNA looping to permit this interaction. Similar to ISWI1, the knockdown of ICOP paralogs caused both IES retention and elevated alternative IES excision (Fig 4, 5). Generally, the levels of alternative excision do not exceed background levels (Singh et al. 2022), but alternative excision is prominent when the ISWI1 complex is disrupted. This led to the emergence of IESs of the “forbidden” peak length. In the *ICOP* KDs, the alternatively excised IESs in the “forbidden” peak mainly originated from the third peak containing longer IESs. This aligns with the observation that partial internal excision, leading to shorter lengths, dominated alternative excision events in *ICOP* KDs (mainly single KDs). In *ISWI1*-KD, partial internal and external excision contributed equally to the alternatively excised IESs and the “forbidden” peak. The difference in excision preference might be caused by ISWI’s ability to move nucleosomes on its own (Längst and Becker 2001; Havas et al. 2000). Some nucleosome repositioning may still happen via ISWI1 in the *ICOP* KDs, although not as effectively as with the ICOPs, leading to easier internal boundary access. However, in *ISWI1*-KD, where nucleosome repositioning fails, IES removal occurs at the next available TA, whether internal or external to the IESs.

In our experiments, nucleosome density differences in *ICOP1/2/PGM*-KD and *ISWI1/PGM*-KD showed sharply peaked distributions, indicating there is not much difference in nucleosome density on IESs in the presence or absence of the ISWI1 complex (Fig 6B). However, *NOWA1/2/PGM*-KD and *PTCAF1/PGM*-KDs showed broader distributions than observed for the ISWI1 complex, implying that the nucleosome densities on IESs are less influenced by the downregulation of chromatin remodeling components than by the downregulation of other genes. Since nucleosome densities do not capture the exact position of the nucleosome, the nucleosome position rather than the number of nucleosomes may change in *ICOP1/2/PGM*-KD and *ISWI1/PGM*-KD. It is challenging to map nucleosome positions precisely in the developing MAC since the DNase sequencing data comprises both old MAC and new MAC sequences.

*NOWA1/2/PGM*-KD and *PTCAF1/PGM*-KDs might have stronger effects on nucleosome density differences because *NOWA1* and *PTCAF1* are expressed earlier in development than the ISWI1 complex and localize to the maternal as well as developing MAC (Nowacki et al. 2005; Ignarski et al. 2014). Therefore, the differences observed in nucleosome densities could either be due to disruption of events downstream of NOWA1 and PTCAF1 functions or due to inter-generational nuclear crosstalk effects on gene regulation as proposed recently (Bazin-Gélis et al. 2023). Irrespective, a clear difference on both chromatin and IES excision can be observed between the ISWI1 complex and other genome editing components, indicating a distinct role for ICOPs and ISWI1 on nucleosomes.

ICOP paralogs might contribute to the directionality of the remodeling complex. In contrast to ISWI1, their knockdown caused a preference, both for partial internal excision (Ext. Fig 4B) and for higher nucleosome densities on long/highly retained IESs (Fig 7B). Higher nucleosome densities might be a direct cause for preferred partial internal excision. We previously proposed a “clothed” model for IES excision, where mispositioned nucleosomes change the accessibility of the IES boundaries to the PGM excision complex (Singh et al. 2022). Assuming that the cooperating PGMs cannot interact across a nucleosome unless a long DNA loop is formed, partial internal excision might be preferred if a nucleosome is located on a TA boundary since an alternative TA lying within the IES might be more easily accessible than a TA outside the IES.

Besides nucleosome positioning, precise targeting of IESs boundaries might also depend on the DNA topology, which influences protein binding and can be exploited as a regulatory mechanism (Baranello et al. 2012). It has been shown that chromatin remodelers of the ISWI family can change the DNA topology (Havas et al. 2000), which might cause the PGM complex to recognize the wrong TA dinucleotides as boundaries if alterations in chromatin remodeling occur. This would also explain how the “forbidden” peak can emerge. According to the original “naked” DNA model, the symmetry of the PGM excision machinery cannot excise 34 - 44 bp fragments (Arnaiz et al. 2012). However, if the DNA helix conformation changes, the PGM complex working distance might correspond to the forbidden length. It seems that the ICOPs can partially compensate for each other since the double KD resembled the *ISWI1*-KD more than the single KDs in terms of cell survival (Fig 3B) and the effects on IES retention (Fig 3D), including alternative excision (Fig 4B, Ext. Fig 4B). We thus propose that the ICOP proteins assist ISWI1’s function in precise genome editing, either by nucleosome sliding or DNA topology changes.

*Paramecium* linker DNA between nucleosomes from the somatic nucleus was shown to be extremely short at just a few bp (Gnan et al. 2022), and no linker histone H1 was detected in *Paramecium* (Drews et al. 2022b). Furthermore, histone modifications characteristic of eu- and heterochromatin in other eukaryotes did not show the expected relations with active and repressive gene expression in *Paramecium* (Drews et al. 2022b). The properties of nucleosomes in *Paramecium* MICs and MACs, including their distribution and dynamics, still need more thorough investigation. Future studies enabling more precise positioning of nucleosomes (esp. via isolation from sufficient flow-sorted MACs) will be essential to determine how nucleosome occupancy and movements, including by the ISWI1 complex, affect the targeting of IESs for excision.

## Materials and methods

### Cultivation of *Paramecium*

Mating type 7 cells (strain 51) of *Paramecium tetraurelia* were grown according to the standard protocol (Beisson et al. 2010c, 2010b). *E. coli* strain HT115 was used for feeding, and the cultures were maintained either at 27 °C or at 18 °C.

### RNAi assay

ICOP1 and ICOP2 RNAi constructs were made by cloning a 538 bp (2708-3246) and a 1089 bp gene fragment (3349-4527), respectively, into the L4440 plasmid. The plasmids were transformed into HT1115 (DE3) *E*. *coli* strain. Knockdown experiments were performed as previously described (Beisson et al. 2010d). Isopropyl ß-D-1-thiogalactopyranoside (IPTG) induction was done at 30 °C. After the cells finished autogamy, 30 post-autogamous cells were fed with a non-induced feeding medium to assay survival. Genomic DNA was extracted from post-autogamous cultures using the standard kit protocol (G1N350, Sigma-Aldrich). PCRs were done on different genomic regions flanking an IES (Supplemental methods Table 1) to test IES retention.

### DNA microinjection and localization

The standard DNA microinjection protocol was followed (Beisson et al. 2010a). Since endogenous regulatory regions failed to express ICOP1 and ICOP2 fusion genes, the regulatory regions of ISWI1 (Singh et al. 2022) were used instead. Human influenza hemagglutinin (HA) was fused N-terminally to ICOP1 and C-terminally to ICOP2. Cells were collected during different stages of autogamy and either stored in 70% ethanol at -20 °C or directly fixed with 2% paraformaldehyde (PFA) in PHEM (PIPES, HEPES, EGTA, Magnesium Sulphate), washed (2 × 5 min at room temperature (RT)) and blocked (1 h at RT) in 5% BSA with 0.1% Triton X-100. Cells were stained overnight at 4 °C with a primary anti-HA antibody (sc-7392, Santa Cruz) followed by washing and secondary anti-mouse Alexa-594 conjugated antibody (BLD-405326, Biozol) incubation for 1 h at RT. After washing, cells were counterstained with DAPI (4,6-diamidino-2-2-phenylindole) in 5% BSA with 0.1% Triton X-100. Cells were mounted with 40 µl of Prolong Gold Antifade mounting medium (Invitrogen). Images were acquired with a Leica SP8 confocal microscope system with a 60× oil objective (NA 1.4). Images were analyzed using Fiji (version 2.9.0/1.53t). Macros used for image analysis are available from https://github.com/Swart-lab/ICOP_code/tree/main/Postprocessing_IF.

### Co-immunoprecipitation and western blot

Co-immunoprecipitation and western blots were done as previously described (Singh et al. 2022). Sonication used an MS72 tip on a Bandelin Sonopulse device with 52% amplitude for 15 s. For non-crosslinked samples, cells were lysed using sonication on ice after washing with 10 mM Tris pH 7.4 in a resuspension of 2 ml lysis buffer. Pulldown fractions were resolved on 12% SDS-PAGE gels. 1% of total lysates were loaded as input, optionally 1% of supernatant after beads incubation as unbound, and 30% (Fig1) or 20% (Ext. Fig 2) of the total IP samples were loaded.

An anti-HA antibody (1:500, sc-7392 HRP, Santa Cruz) and anti-GFP antibody (1:2000, ab290, Abcam) incubation was done overnight at 4 °C. The secondary antibody, goat-anti-Rabbit HRP conjugated (12-348, Merck Millipore), was incubated for 1 h at room temperature. Membranes were screened using AI600 (GE Healthcare).

### Plasmids and vectors for recombinant protein expression assay

DNA sequences coding for *Paramecium* proteins ISWI1, ICOP1, and ICOP2 were codon-optimized (Supplemental methods Table 5) for expression in *E. coli* using the GENEius tool of Eurofins (Luxembourg). Gene synthesis was performed at Eurofins Genomics Germany GmbH (Ebersberg, Germany). The synthetic constructs were cloned into pET-MCN vectors (Romier et al. 2006), expressing proteins with either no tag, a hexahistidine (His), or a GST tag. Codon-optimized sequences are provided with Supplemental methods. Plasmids were co-transformed in different combinations into *E. coli* strain Gold pLysS.

### Protein expression in *E*. *coli*

100 µl of LB culture was added to 50 ml of ZY medium (Studier 2014) containing appropriate antibiotics. Cultures were grown at 37 °C at 180 rpm until an OD 600 of 2 was reached. Afterward, the cultures were incubated at 20 °C at 180 rpm overnight for protein expression. After overexpression, 2 ml of the culture was centrifuged at 4000 g at 4 °C, and the cell pellets were frozen at -80 °C.

### Co-precipitation of recombinant proteins

Cell pellets were resuspended in 1 ml of lysis buffer: 20 mM Tris pH 7.5, 100 mM NaCl for GST pulldown or 20 mM Tris pH 7.5, 100 mM NaCl, 20mM Imidazole, 1mM DTT for His pulldown. 20% amplitude (0.5 s on, 0.5 s off) with an MS72 tip (Bandelin Sonopulse) was used for sonication, followed by centrifugation (21130 g, 15 min, 4 °C) to recover the supernatant for pulldown. 30 µl of beads (42172.01/ 42318.01, Serva) were washed once with 1 ml of Milli-Q water to remove ethanol and centrifuged (2 min at 1000 g at 4 °C, also for subsequent bead centrifugation steps). Beads were equilibrated using 1 ml of lysis buffer and centrifuged once. The supernatant was incubated with the beads for 1 h or overnight at 4 °C using gentle shaking. After three washes, beads were resuspended into 30 µl of 2× protein loading Buffer (100 mM Tris-HCl pH 6.8, 4% (w/v) SDS, 20% Glycerol, 0.2 M DTT), boiled for 10 min, and centrifuged briefly before loading supernatant on a 10-12% SDS-PAGE gel. 1% of the total lysate was loaded as input, and 20% of the total pulldown was loaded in the IP fraction. 1:4000 rabbit anti-GST antibody (G7781, Sigma) and mouse anti-His (1:2500, 362601, BioLegend) were diluted in 5% BSA in 1× PBS + 0.2% Tween20 for blotting. 1:5000 reciprocal secondary antibody incubation was done for 1 h at room temperature. Membranes were screened on an AI600 (GE Healthcare).

### DNA and total RNA extraction and sequencing

Standard methods were used to isolate macronuclear DNA and total RNA for sequencing. Detailed protocols are provided in the Supplemental methods.

### IES retention and alternative boundary analysis

Whole genome sequencing (WGS) reads of enriched new MAC DNA after knockdown were trimmed for Illumina adapter sequences using TrimGalore (Krueger 2019) (Supplemental Materials and Methods Table 2). ParTIES (Denby Wilkes et al. 2016) v1.05 was used to map reads to MAC and MAC+IES genomes and calculate IRSs. To accommodate changes in a newer version of samtools (Li et al. 2009), the /lib/PARTIES/Map.pm file was changed (Supplemental methods Table 3). IRSs are provided in SourceData_Fig4 (Singh 2023) as ICOP_IRS.tab.gz. IRS correlations using IRSs form published knockdown data ((*ISWI1*-KD (Singh et al. 2022), *PGM*-KD (Arnaiz et al. 2012), *TFIIS4*-KD (Maliszewska-Olejniczak et al. 2015) and *PTIWI01/09*-KD (Furrer et al. 2017)) were calculated with After_ParTIES (option -- use_pearson (https://github.com/gh-ecs/After_ParTIES)).

Since alternative excision analysis depends on IES coverage, to ensure a fair comparison, libraries were adjusted to similar sizes by downsampling. Downsampling factors relative to the smallest library used were calculated according to the number of properly paired and mapped reads to the MAC+IES reference genome (ND7 = 0.686; ICOP1 = 0.512; ICOP2 = 0.453; ISWI1 = 0.698; ICOP1_2 = 1.0). The “MILORD” module of a ParTIES pre-release version (13 August 2015) was used to annotate alternative and cryptic IES excision (SourceData_Fig5; (Singh 2023)).

Reference genomes used for these analyses are indicated in Supplemental methods. All scripts are available from https://github.com/Swart-lab/ICOP_code/tree/main/Alternative_excision.

### Nucleosomal DNA Isolation and Illumina DNA-sequencing

Nucleosomal DNA was isolated with the EZ Nucleosomal DNA Prep Kit (D5220, Zymo Research) as previously described (Singh et al. 2022), except that digested DNA was size-selected with SPRIselect magnetic beads (Beckman Coulter) to enrich for mono- and di-nucleosomal fragments (0.7× volume right-side size selection). Libraries were prepared with NEBNext Ultra II DNA library prep kit (E7645S, NEB), size-selected for 150 bp insert. 2×100 bp paired-end sequencing was performed on an Illumina NextSeq 2000 instrument with P3 chemistry at MPI for Biology, Tübingen.

### Nucleosome Density Analysis

Illumina adapter sequences were trimmed from reads with TrimGalore (Krueger 2019) (Supplemental Materials and Methods Table 2). Nucleosome densities were acquired as previously described (Singh et al. 2022). Reads were mapped to the MAC+IES genome, then properly paired and mapped reads overlapping IESs were extracted and counted. DNase reads were size selected (100 - 175 bp outer distance). Library sizes to calculate downsampling factors were retrieved with the “samtools stats” command on the .sorted.bam files. The length distribution of outer distances of PE reads mapping to scaffold51_9 was plotted (Ext. Fig 5B).

Samples used for nucleosome density analysis are provided in Supplemental methods (Table 6). Nucleosome density differences (re_rc) were calculated for each IES by subtracting the nucleosome density of the control (r_c) from the experimental sample (r_e).

re_rc = r_e - r_c

IES with infinite (“inf”) or not available “nan” values were excluded, resulting in 43,409 (in *NOWA1/2/PGM*-KD) and 44,448 (in *ICOP1/2/PGM*-KD) IESs used for analysis. Kolmogorov-Smirnov (KS) statistics and associated p-values for two sample tests were calculated to assess distribution differences.

All scripts are available from https://github.com/Swart-lab/ICOP_code/tree/main/Nucleosome_density.

Read counts on IESs are available in SourceData_Fig7 (Singh 2023).

### sRNA analysis

sRNA-seq was mapped to the *Paramecium tetraurelia* strain 51 MAC + IES genome and L4440 silencing vector with bwa version 0.7.17-r1188 (Li and Durbin 2009). 10-49 bp long, uniquely mapped reads (possessing the flags “XT:A:U”) were selected by grep in a shell script. sRNA length histograms were generated by a Python script. Shell scripts for the RNA mapping, post-processing, and histogram are available from https://github.com/Swart-lab/ICOP_code/tree/main/sRNA_analysis.

### Knockdown efficiency validation using RNA-seq

Total RNA was sequenced by Genewiz (Germany, GmbH) using poly-A enrichment with NovaSeq 2×150 bp reads. Illumina adapter sequences were trimmed from reads with TrimGalore (Krueger 2019) (Supplemental Materials and Methods Table 2). Reads were mapped to the *Paramecium tetraurelia* strain 51 transcriptome (Supplemental methods). Mapping showed high coverage on the silencing regions, most likely caused by RNAs of the siRNA silencing pathway. For each knockdown, target gene was replaced by three split transcripts (the silencing region, the 5’ upstream non-silencing region and the 3’ downstream non-silencing region), and only the 5’ upstream region was considered for analysis. FPKM (fragments per kilobase transcript per million mapped reads) values were calculated using eXpress (Roberts and Pachter 2013) (SourceData_Fig4; (Singh 2023)) and rounded by the standard Python method to integers. Scripts are available from https://github.com/Swart-lab/ICOP_code/tree/main/KD-efficiency.

### Structure prediction with AlphaFold

Protein structures were predicted with AlphaFold multimer version 2.2.0 and 2.3.0 (Evans et al. 2021; Jumper et al. 2021). Protein sequences provided as input are listed in the Supplemental methods (Table 4). All predictions were computed on the high-performance computer “Raven”, operated by the Max-Planck Computing and Data Facility in Garching, Munich, Germany. PDB files are available as SourceData_Fig3 (Singh 2023).

## Competing Interest Statement

The authors declare no competing interests.

## Acknowledgments

We thank the MPI for Biology (Tübingen, Germany) core facilities for microscopy and sequencing assistance; Vikram Alva for helpful discussions on using HHpred; A. Noll for computer system administration.

## Author contributions

A.S., L.H., E.C.S. designed research; A.S., L.H., C.E., E.N., B.K.B.S., performed research; A.S., L.H., E.C.S. analyzed data; M.N., E.C.S. contributed reagents/analytical tools; L.H., A.S., E.C.S., wrote the paper; A.S., F.B., E.C.S supervision.

## Data availability

EDMOND: https://doi.org/10.17617/3.ZBOLU8

ENA: PRJEB64685

ProteomExchange: PXD044340

---

**Extended Figure 1:**
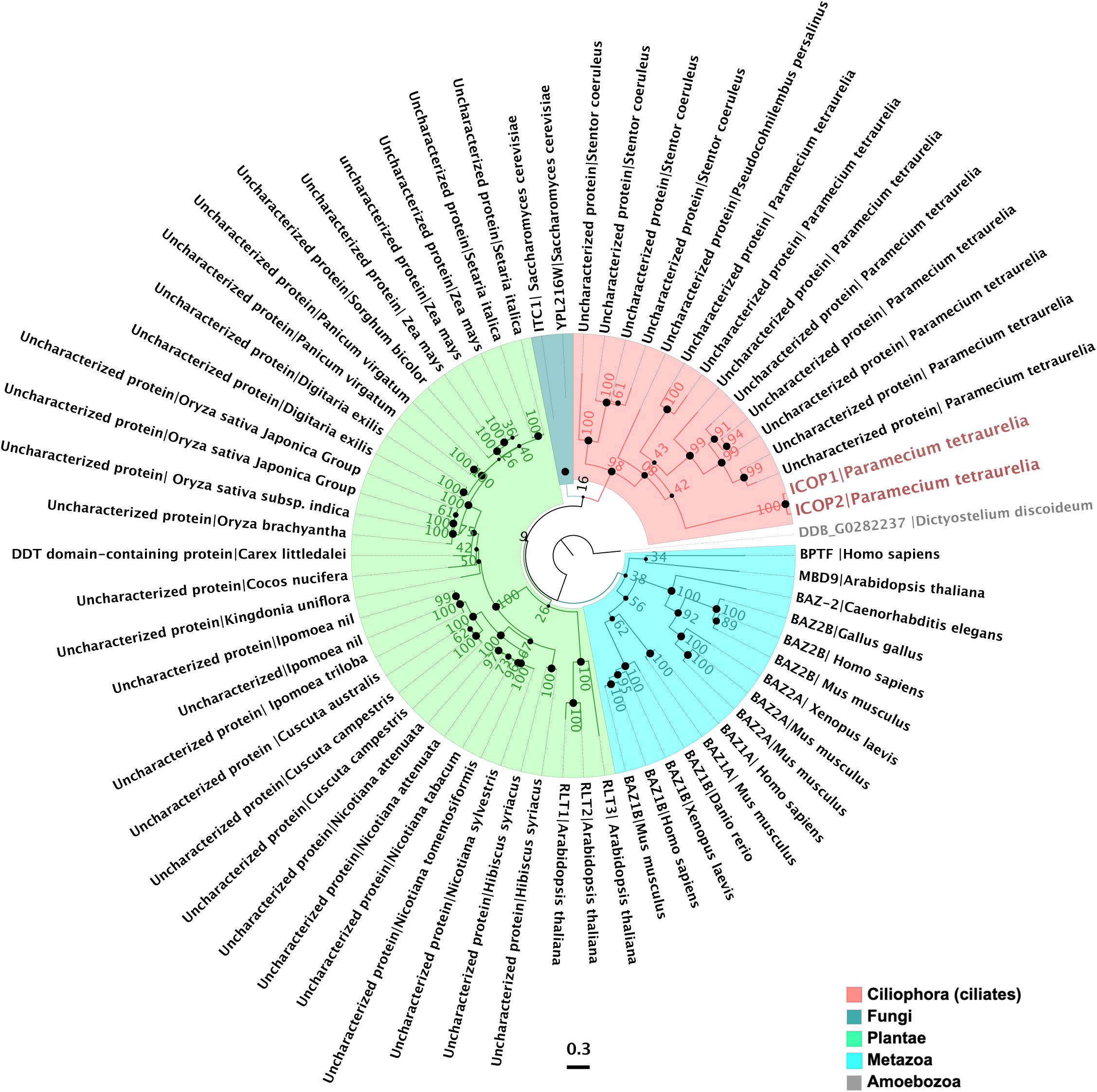
WSD-containing proteins are highly diverse. Phylogenetic analysis of proteins with WHIM2 domain in selected organisms. Node bootstrap values are labeled, and the ’•’ size corresponds to node values. The tree is rooted at *Dictyostelium discoideum*, labeled in gray. Scale bar is 0.3. ICOP1 and ICOP2 are labeled in salmon

**Extended Figure 2:**
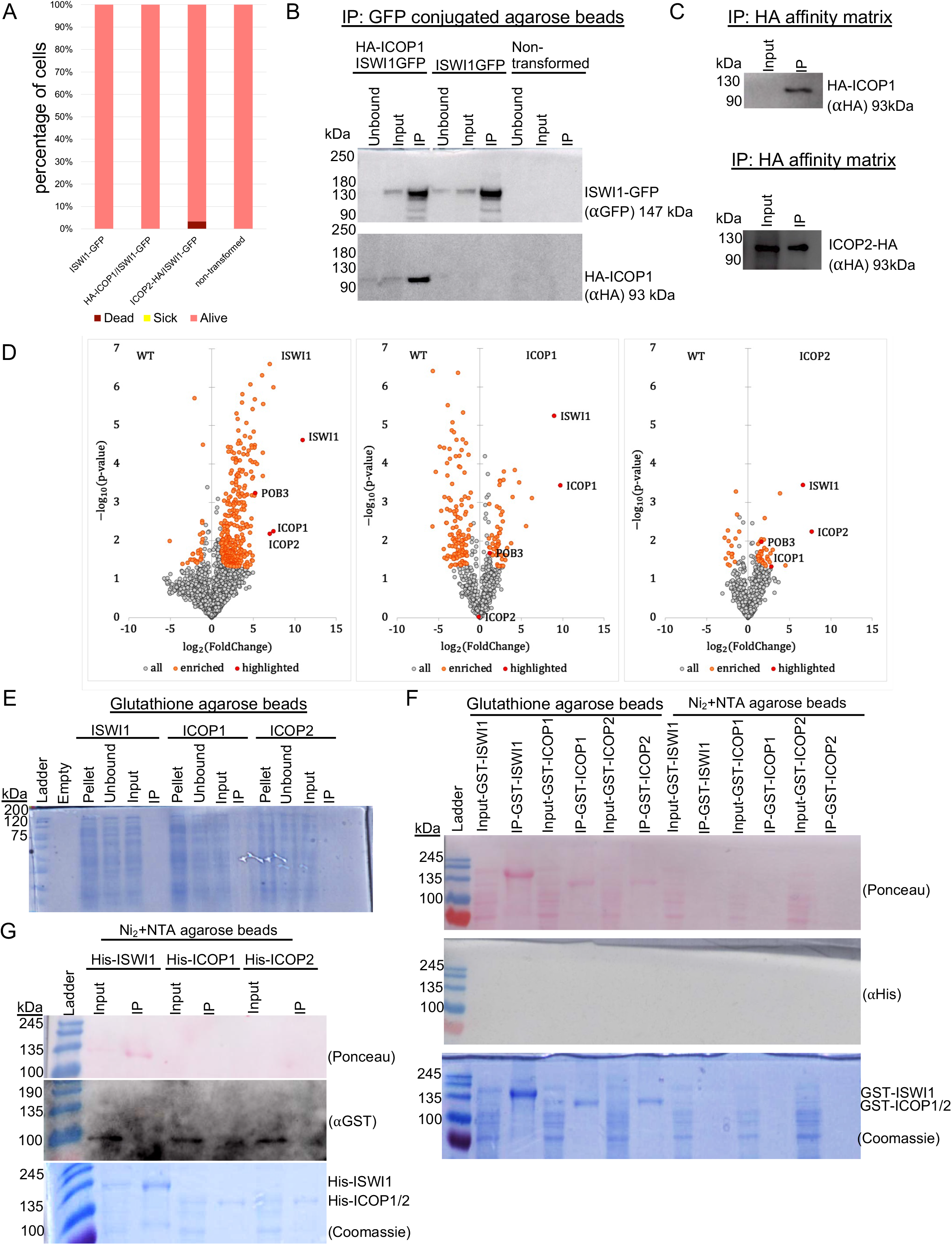
ICOP paralogs interact with ISWI1. (A) Survival assay on F1 generation after knockdown. Alive (pink): normal division. Sick (yellow): slower division rate. Dead (Cayenne): no cells. (B) Western blot on co-IP of HA-ICOP1/ISWI1-GFP co-transformed, ISWI1-GFP transformed and non-transformed, wild-type *Paramecium*. (C) Western blot on co-IP of HA-ICOP1 and ICOP2-HA overexpressed in paramecia. (D) Volcano plots showing protein enrichment of mass spectrometry (MS) analysis for ISWI1-GFP (left), HA-ICOP1 (middle), and ICOP2-HA (right) co-IP. (E) to (F): Pulldowns on overexpressed recombinant proteins in *E. coli*. (E) Coomassie staining of untagged ISWI1, ICOP1 and ICOP2. (F) Western blot and Coomassie staining of GST-tagged recombinant protein pulldowns; Ponceau-stained membranes probed with anti-His antibody. (G) Western blot and Coomassie staining of His-tagged recombinant proteins; Ponceau-stained membranes probed with anti-GST antibody.

**Extended Figure 3:**
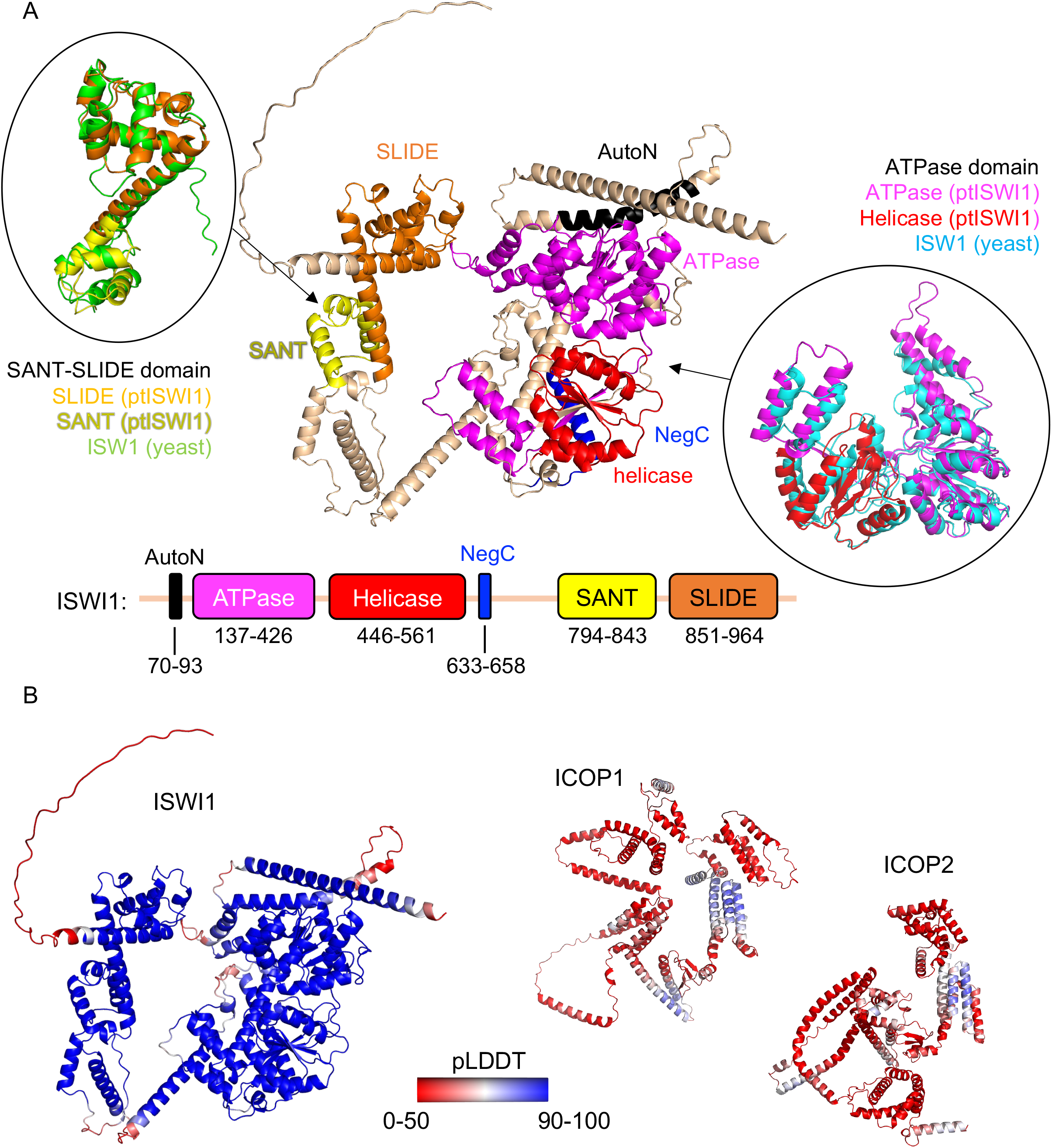
ISWI1 and ICOP structure predictions. (A) and (B) AlphaFold (version 2.2.0) structure predictions. (A) Domains in *Paramecium* ISWI1. ATPase and Helicase are superimposed with a published structure of N-terminal ISWI from yeast (PDB accession number 6JYL) (color: cyan) and SANT-SLIDE domains are superimposed with ISW1a (del_ATPase) from yeast (PDB accession number 2Y9Y) (color: green). (B) Structure prediction confidence for ISWI1, ICOP1, and ICOP2. Models are colored by predicted local distance difference test (pLDDT). pLDDT ≤ 50 are represented in red. pLDDT ≥ 90 are represented in blue.

**Extended Figure 4:**
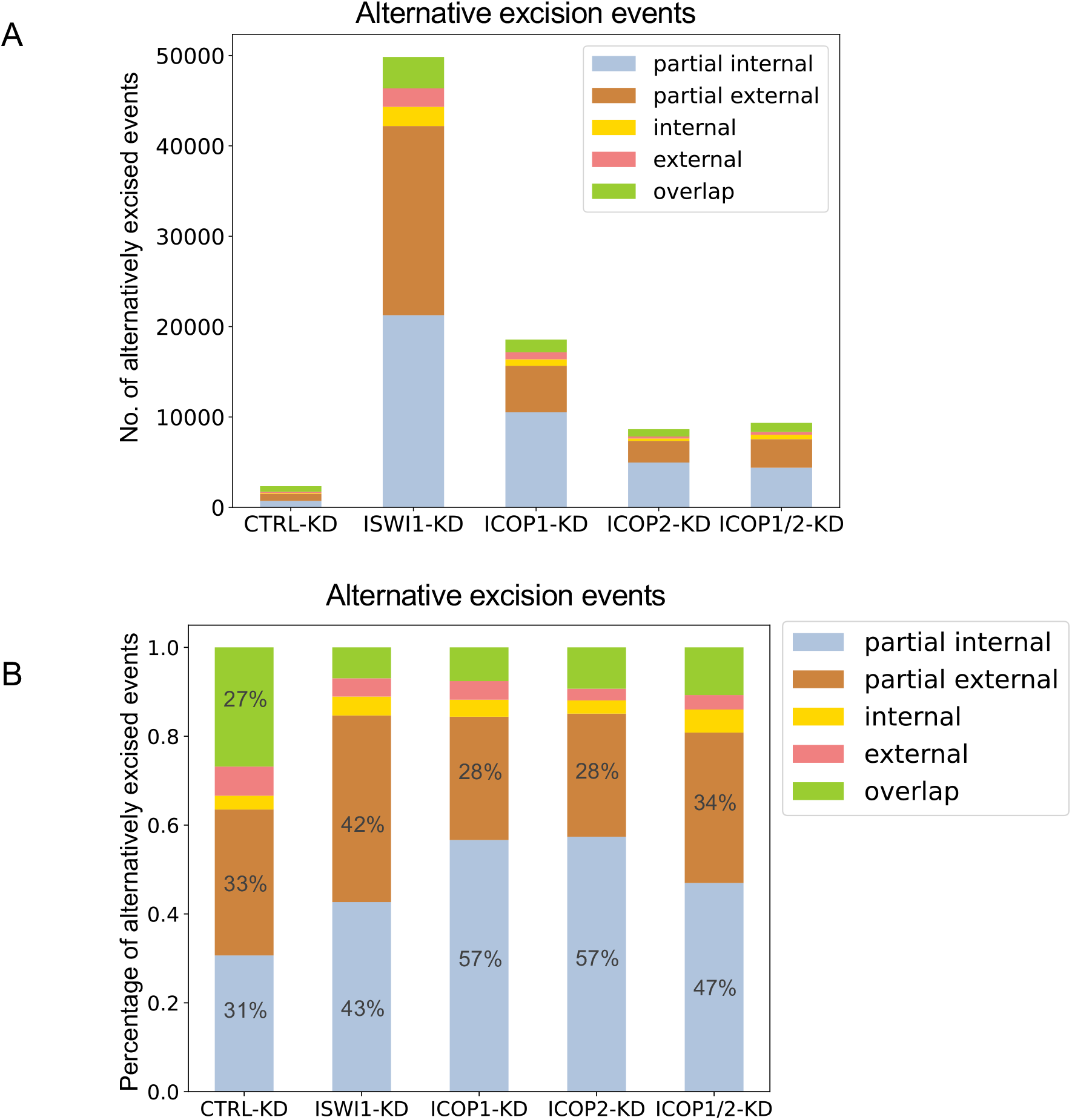
Alternative excision events. Stacked bar graphs of alternative excision events detected in *ISWI1*-KD, *ICOP1*-KD, *ICOP2*-KD and *ICOP1/2*-KD. *ND7*-KD was used as a control. (A) Absolute and (B) relative abundance of alternative excision events occurring upon KDs.

**Extended Figure 5:**
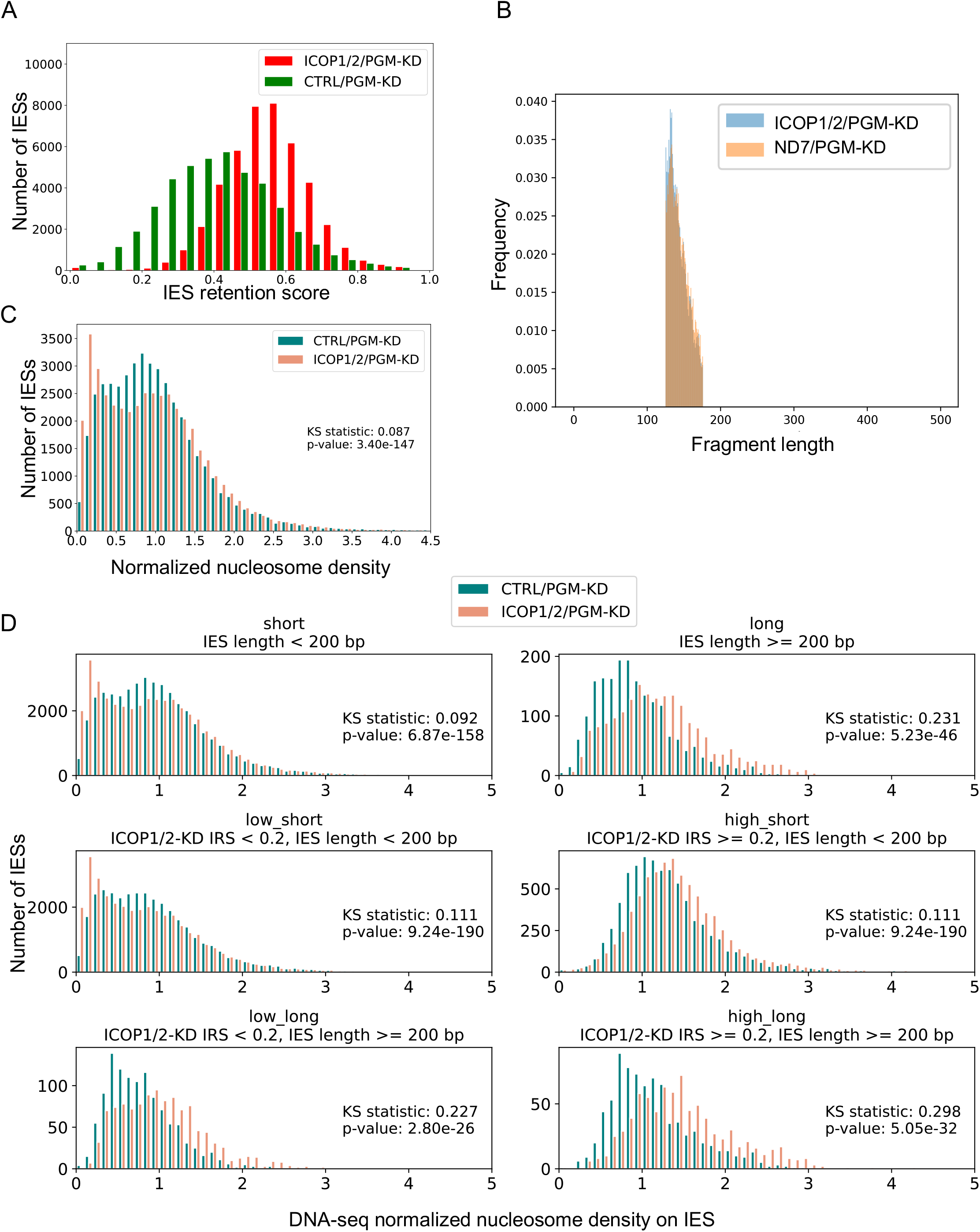
Nucleosome densities for *ICOP1/2/PGM*-KD and *CTRL/PGM*-KD. (A) IRS histogram for *ICOP1/2/PGM*-KD and *CTRL/PGM*-KD. (B) Size distribution of reads on scaffold51_9 for *ICOP1/2/PGM*-KD and *CTRL/PGM*-KD. (C) Nucleosome densities on all IESs in *ICOP1/2/PGM*-KD and *CTRL/PGM*-KD. (D) Nucleosome densities on selected IES groups in *ICOP1/2/PGM*-KD and *CTRL/PGM*-KD. IESs were grouped by IES retention score (IRS) in *ICOP1/2*-KD (low: IRS < 0.2; high: IRS ≥ 0.2) and IES length (short: IES length < 200 bp; long: IES length ≥ 200 bp). IES group is given above the diagrams.

**Extended Figure 6:**
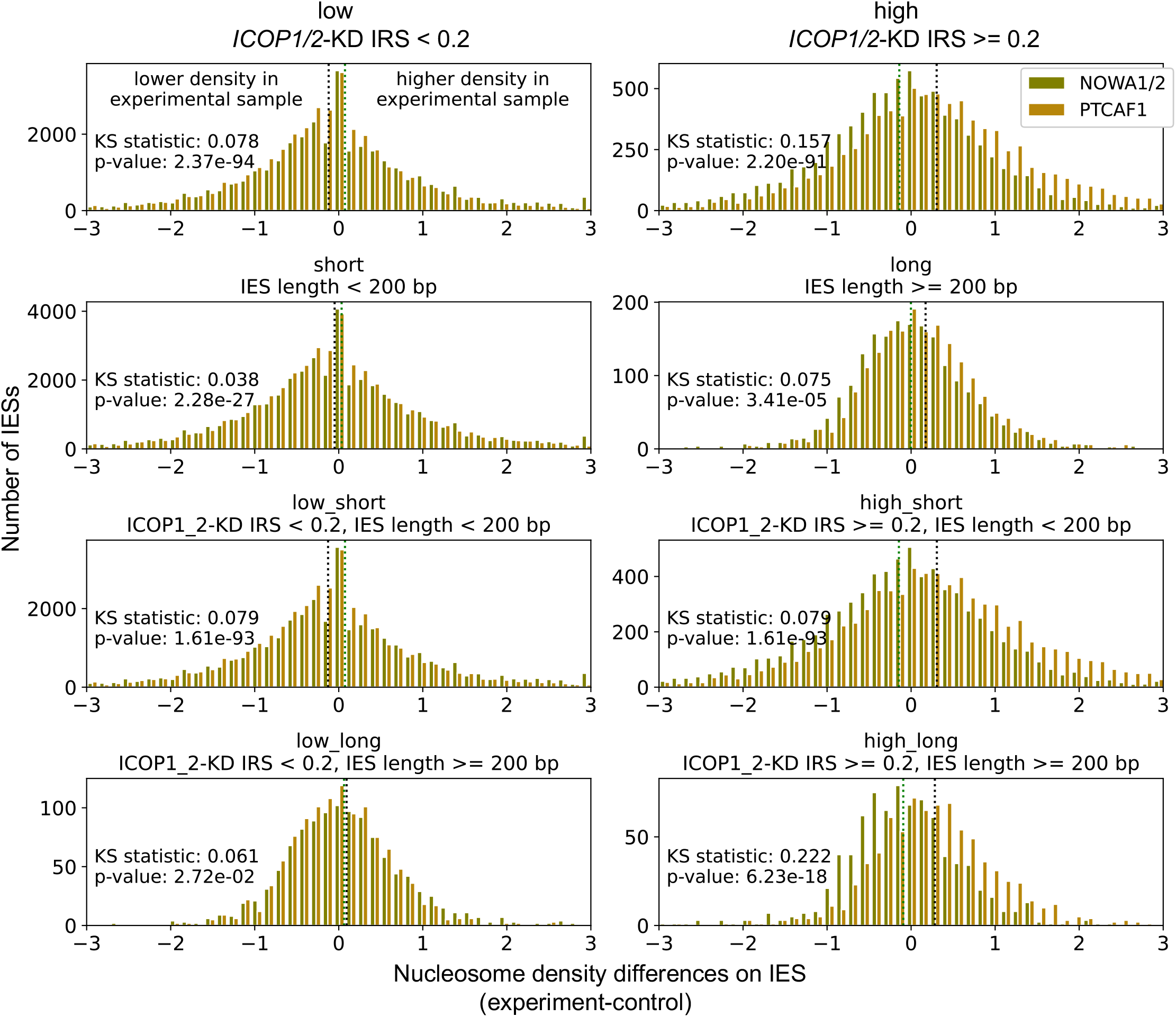
Nucleosome density differences for *NOWA1/2/PGM*-KD and *PTCAF1/PGM*-KD. Comparison of *NOWA1/2/PGM*-KD and *PTCAF1/PGM*-KD nucleosome density differences in selected IES groups: IESs were grouped by IES retention score (IRS) in *ICOP1/2*-KD (low: IRS < 0.2; high: IRS ≥ 0.2) and IES length (short: IES length < 200 bp; long: IES length ≥ 200 bp). The specification for each IES group is given above the individual diagrams. Means as dashed lines (*NOWA1/2/PGM*-KD: green; *PTCAF1/PGM*-KD: black).

**Extended Table 1:** Predicted interactions in AlphaFold2 models.

| **AF2 version** | **model** | **interaction predicted** | **GxD involved** | **interaction interface** |
| --- | --- | --- | --- | --- |
| v2.2.0 | ISWIN + ICOP1 | yes | no | ICOP1 resi 556-597; ISWI1 resi 425-426+431+434-437+474+477-478+481 |
| v2.2.0 | ISWIN + ICOP2 | yes | no | ICOP2 resi 560-603; ISWI1 resi 425-426+431+434-437+474+477-478+481 |
| v2.2.0 | ISWIC + ICOP1 | no | - | - |
| v2.2.0 | ISWIC + ICOP2 | no | - | - |
| v2.2.0 | ISWI1 + ICOP1 | no | - | - |
| v2.2.0 | ISWI1 + ICOP2 | no | - | - |
| v2.2.0 | ICOP1 + ICOP2 | no | - | - |
| v2.2.0 | ISWIN + ICOP1/2 | no | - | - |
| v2.2.0 | ISWIN + Ptiwi01 | no | - | - |
| v2.2.0 | ISWIC + Ptiwi01 | yes | - | - |
| v2.3.0 | ISWIN + ICOP1 | yes | no | large interaction interface |
| v2.3.0 | ISWIN + ICOP2 | yes | no | large interaction interface |
| v2.3.0 | ISWIC + ICOP1 | yes | no | large interaction interface |
| v2.3.0 | ISWIC + ICOP2 | yes | no | large interaction interface |
| v2.3.0 | ISWI1 + ICOP1 | yes | no | mostly ISWI1 N-terminus |
| v2.3.0 | ISWI1 + ICOP2 | yes | no | mostly ISWI1 N-terminus |
| v2.3.0 | ICOP1 + ICOP2 | yes | - | - |

**Extended Table 2:** Percentage of alternatively excised IESs.

|  | including IES with 100%<br>alternative excision |  |  | excluding IES with 100%<br>alternative excision |  |  |
| --- | --- | --- | --- | --- | --- | --- |
|  | median | mean | IESs | median | mean | IESs |
| *CTRL*-KD | 0.00 % | 2.59 % | 41311 | 0.00 % | 1.09 % | 40680 |
| *ISWI1*-KD | 4.55 % | 10.86 % | 43983 | 4.35 % | 9.18 % | 43171 |
| *ICOP1*-KD | 0.00 % | 8.97 % | 42237 | 0.00 % | 7.01 % | 41349 |
| *ICOP2*-KD | 0.00 % | 6.00 % | 41573 | 0.00 % | 4.18 % | 40785 |
| *ICOP1/2*-KD | 0.00 % | 6.53 % | 41767 | 0.00 % | 4.71 % | 40972 |

Median and mean percentage (in %) of alternative excision for all IESs in the KDs.
The number of IESs included in the analysis is given ("IESs"). IESs with 100% alternative excision are either included (left) or excluded (right).

**Extended Table 3:** Lengths of alternative excision.

|  | **minimum** | **maximum** | **mean** | **median** |
| --- | --- | --- | --- | --- |
| ref_length | 20 | 5314 | 79.13 | 50 |
| *CTRL*-KD | 9 | 9032 | 260.55 | 77 |
| *ISWI*1-KD | 5 | 9892 | 208.64 | 73 |
| *ICOP*1-KD | 6 | 9590 | 197.73 | 67 |
| *ICOP*2-KD | 5 | 9467 | 143.04 | 65 |
| *ICOP*1/2-KD | 5 | 9878 | 185.16 | 67 |

Length of excised fragments in IES excision for different KDs. ref\_length: Reference IES lengths in standard IES excision.

**Extended Table 4:**
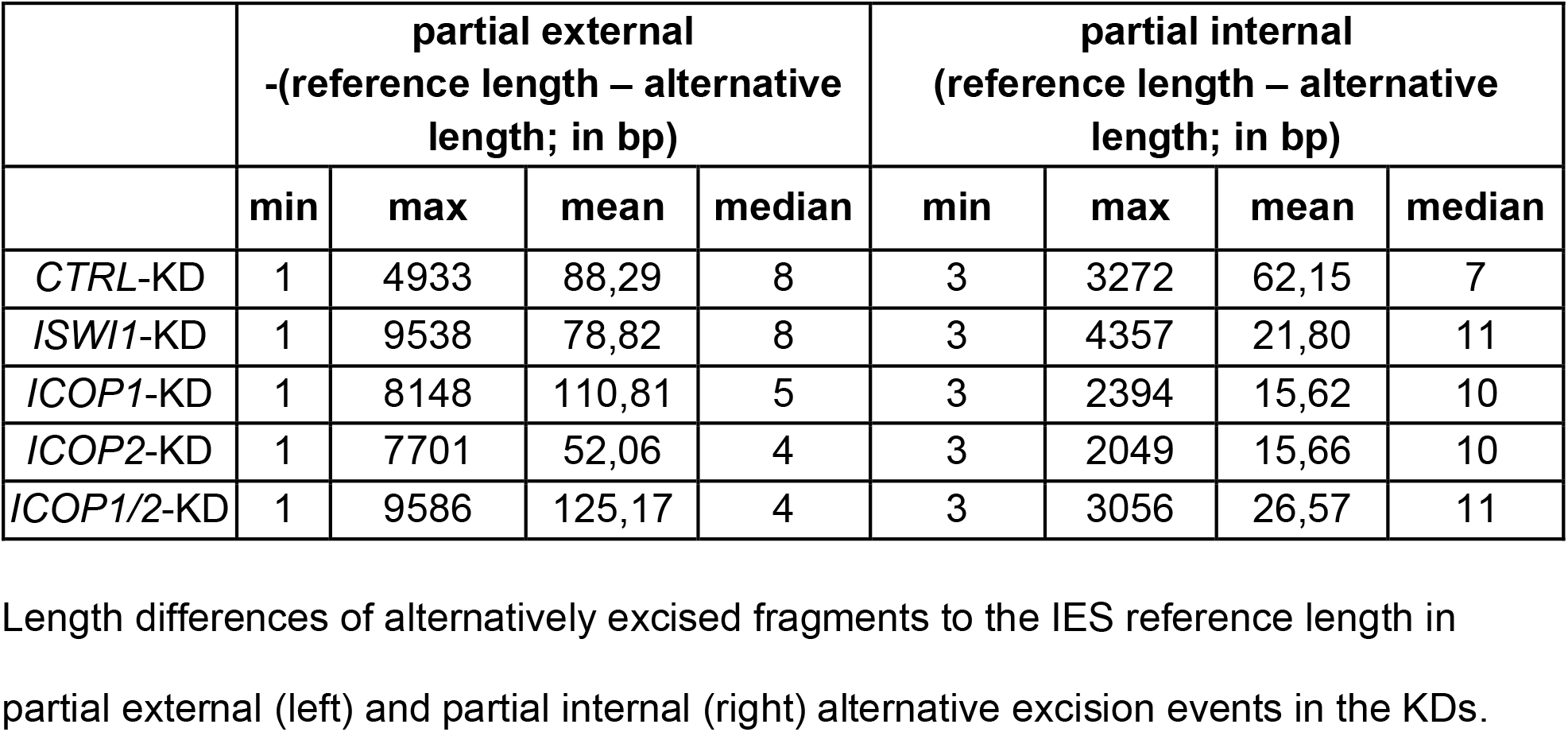
Length differences in partial external and partial internal alternative excision.

|  | partial external<br>-(reference length – alternative<br>length; in bp) |  |  |  | partial internal<br>(reference length – alternative<br>length; in bp) |  |  |  |
| --- | --- | --- | --- | --- | --- | --- | --- | --- |
|  | min | max | mean | median | min | max | mean | median |
| *CTRL*-KD | 1 | 4933 | 88,29 | 8 | 3 | 3272 | 62,15 | 7 |
| *ISWI1*-KD | 1 | 9538 | 78,82 | 8 | 3 | 4357 | 21,80 | 11 |
| *ICOP1*-KD | 1 | 8148 | 110,81 | 5 | 3 | 2394 | 15,62 | 10 |
| *ICOP2*-KD | 1 | 7701 | 52,06 | 4 | 3 | 2049 | 15,66 | 10 |
| *ICOP1/2*-KD | 1 | 9586 | 125,17 | 4 | 3 | 3056 | 26,57 | 11 |

Length differences of alternatively excised fragments to the IES reference length in partial external (left) and partial internal (right) alternative excision events in the KDs.

**Extended Table 5:** Means of nucleosome density differences.

| IES group | IRS in *ICOP1/2*-KD | IES length | *ICOP1/2* | *ISWI1* | *NOWA1/2* | *PTCAF1* |
| --- | --- | --- | --- | --- | --- | --- |
| total | all | all | -0.03 ± 0.78 | -0.02 ± 1.17 | 0.03 ± 1.69 | -0.04 ± 1.45 |
| low | < 0.2 | all | -0.07 ± 0.77 | 0.00 ± 1.07 | 0.07 ± 1.75 | -0.12 ± 1.43 |
| high | ≥ 0.2 | all | 0.15 ± 0.81 | -0.09 ± 1.50 | -0.14 ± 1.40 | 0.30 ± 1.48 |
| short | all | < 200 bp | -0.04 ± 0.79 | -0.01 ± 1.02 | 0.03 ± 1.72 | -0.05 ± 1.44 |
| long | all | ≥ 200 bp | 0.29 ± 0.39 | -0.09 ± 2.88 | 0.00 ± 0.92 | 0.17 ± 1.75 |
| low_short | < 0.2 | < 200 bp | -0.08 ± 0.78 | 0.01 ± 0.86 | 0.07 ± 1.77 | -0.13 ± 1.40 |
| low_long | < 0.2 | ≥ 200 bp | 0.24 ± 0.38 | -0.15 ± 3.74 | 0.06 ± 1.04 | 0.09 ± 2.19 |
| high_short | ≥ 0.2 | < 200 bp | 0.13 ± 0.84 | -0.10 ± 1.57 | -0.15 ± 1.46 | 0.31 ± 1.54 |
| high_long | ≥ 0.2 | ≥ 200 bp | 0.35 ± 0.41 | -0.01 ± 0.49 | -0.09 ± 0.71 | 0.28 ± 0.78 |

Means and standard deviation are given for the different IES groups in different knockdowns (*ICOP1/2*: *ICOP1/2*/PGM-KD; *ISWI1*: *ISWI1*/PGM-KD; *NOWA1/2*: *NOWA1/2*/PGM-KD; *PTCAF1*: *PTCAF1*/PGM-KD). The characteristics for IESs in each group are specified in "IRS in *ICOP1/2*-KD" and "IES length".

## Notes

https://doi.org/10.17617/3.ZBOLU8

https://github.com/Swart-lab/ICOP_code

### Competing Interest Statement

The authors have declared no competing interest.

