## Supplemental methods for "ISWI1 complex proteins facilitate developmental genome editing in *Paramecium*"

#### Supplemental Materials and Methods

#### HHpred identification of protein domains

ICOP1 (XM_001447768.1, PTET.51.1.P0440186) and ICOP2 (XM_001437312.1, XM_001437313.1, PTET.51.1.P0180124) protein sequences were analyzed using HHpred from the MPI bioinformatics toolkit (https://toolkit.tuebingen.mpg.de/#/) across the COG (Tatusov et al. 2000), Pfam (Finn et al. 2016, 2003), NCBI Conserved Domain Database (CDD) (Marchler-Bauer et al. 2017), and ECOD (Cheng et al. 2014, 2015) databases. PSI-BLAST (Bhagwat and Aravind 2007) with 4 iterations was used to identify further proteins with WSD domains, and multiple alignments were done using MAFFT (Katoh and Standley 2013; Kuraku et al. 2013) provided as a plugin within Geneious (version 2022.1.1, http://www.geneious.com) (Kearse et al. 2012). InterProScan (Paysan-Lafosse et al. 2023) within Geneious was used to identify domains within MAFFT-aligned protein sequences.

#### Phylogenetic analysis of ICOP1 and ICOP2

Trimal-auto (Capella-Gutiérrez et al. 2009) was used to select well-aligned columns from the MAFFT-aligned protein sequences. PHYML version 2.2.4(Guindon and Gascuel 2003) provided as a plugin in Geneious (version 2022.1.1, http://www.geneious.com) was used to generate a maximum likelihood phylogeny with 100 bootstrap replicates. FigTree v1.4.4 (http://tree.bio.ed.ac.uk/software/figtree/) was used to inspect, manipulate and generate the graphical tree.

#### Mass spectrometry analysis

Samples were separated on a 4%–12% NOVEX NuPage gradient gel (Thermo) for 10 minutes at 180 V in 1 X MES buffer (Thermo). Proteins were fixed and stained with Coomassie G250 brilliant blue (Carl Roth). The gel lanes were cut, and each lane was minced into approximately 1x1 mm pieces. Gel pieces were destained with a 50% ethanol/50 mM ammonium bicarbonate (ABC) solution. Proteins were reduced in 10 mM DTT (Sigma-Aldrich) for 1 hour at 56 °C and then alkylated with 50 mM iodoacetamide (Sigma-Aldrich) for 45 min at room temperature. Proteins were digested with 1 µg mass spectrometry grade trypsin (Sigma) overnight at 37 °C. Peptides were extracted from the gel by two incubations with 30% ABC/acetonitrile and three subsequent incubations with pure acetonitrile. The acetonitrile was subsequently evaporated in a concentrator (Eppendorf) and loaded on StageTips (Rappsilber et al. 2007) for desalting and storage.

Peptides were eluted from the StageTips using 80% acetonitrile / 0.1% formic acid and concentrated before loading on an uHPLC nLC-1200 system coupled to an Exploris 480 mass spectrometer (Thermo). The peptides were loaded on a 50 cm (Exploris 480) column (75 µm inner diameter) in-house packed with Reprosil C18 (Dr. Maisch GmbH) and eluted with a 73 or 88 min optimized gradient increasing from 3% to 40% mixture of 80% acetonitrile/0.1% formic acid at a flow rate of 225 nl/min or 250 nl/min. The Exploris 480 was operated in positive ion mode with a data-dependent acquisition strategy of one MS full scan (scan range 300 - 1,650 m/z; 60,000 resolution; normalized AGC target 300%; max IT 28 ms) and up to twenty MS/MS scans (15,000 resolution; AGC target 100%, max IT 40 ms; isolation window 1.4 m/z) with peptide match preferred using HCD fragmentation.

MS raw data were searched using the Andromeda search engine (Cox et al. 2011) integrated into MaxQuant suite 1.6.5.0 (Cox and Mann 2008) using the *Paramecium* predicted proteins as the database (ptetraurelia_mac_51_annotation_v2.0). In all analyses, carbamidomethylation at cysteine was set as a fixed modification, while methionine oxidation and protein N-acetylation were considered as variable modifications. The match between run option was activated. Prior to bioinformatics analysis, reverse hits, proteins only identified by site, protein groups based on one unique peptide, and known contaminants were removed.

For further bioinformatics analysis, the label-free quantitation (LFQ) values were $\log_2$ transformed, and the median across the replicates was calculated. This enrichment was plotted against the – $\log_{10}$ transformed p-value (Welch t-test) using the ggplot2 package in the R environment.

#### Total RNA extraction and mRNA sequencing.

Approximately $1.2 \times 10^6$ cells were collected from the early (approx. 40% of cells with a fragmented MAC) and the late (majority of cells with visible anlagen) developmental stages using an oil centrifuge at 280 g for 2 minutes. The cells were washed twice in 10 mM Tris-HCl (pH 7.4), flash frozen using liquid nitrogen dropped gently over the pellets, and stored at -80°C. Total RNA extraction was performed by adding 6 ml of Tri reagent (Sigma-Aldrich, T9424) per sample and following the standard protocol provided with the reagent. mRNA and sRNA libraries were prepared and sequenced at Genewiz International (Leipzig, Germany).

#### Macronuclear isolation and Illumina DNA-sequencing

Samples for MAC isolation were collected from *ICOP1*-KD*, ICOP2*-KD*,* and *ICOP1/2*-KD cultures three days post autogamy as described previously (Arnaiz et al. 2012). *ICOP1/2/PGM*-KD and *ND7/PGM*-KD cultures were collected ca. 12 h after developing MACs were visible in most of the cells. DNA libraries were prepared using the FS DNA Library Prep kit (E7805, NEB). Paired-end (2×150 bp) sequencing was done on NextSeq 2000 at MPI for Biology, Tübingen.

#### Reference genomes and predicted genes

Reference genome to analyze DNA-seq data:

MAC: https://paramecium.i2bc.paris-saclay.fr/files/Paramecium/tetraurelia/51/sequences/ptetraurelia_mac_51.fa

MAC+IES: https://paramecium.i2bc.paris-saclay.fr/files/Paramecium/tetraurelia/51/sequences/ptetraurelia_mac_51_with_ies.fa

Reference CDS + UTR sequences used to analyze mRNA-seq data:

https://paramecium.i2bc.paris-saclay.fr/files/Paramecium/tetraurelia/51/annotations/ptetraurelia_mac_51/ptetraurelia_mac_51_annotation_v2.0.transcript.fa

### Table 1: Primers used for IES Retention PCR analysis

| IES | Primer sequence (5’ to 3’ orientation) |
|---|---|
| MT Locus F | GGTGTTTATATCTTAATTGTTGACCCTCAC |
| MT Locus R | CCATCTATACTCCATTCTTTATCTTAATTCAT |
| 51A2591F | ATGTGTTTGGACTGGATTGGCATGTAGAAG |
| 51A2591R | GATGTAGCATAACATTTATCAACAATCCAT |
| 51A6649F | ACTGCACCTCTAACTTTAACAAGCGAAGCA |
| 51A6649R | CAGCAGTACATCCAGCTCTCTAAGTTTAGC |

### Table 2: Reads used for adapter trimming

| | |
|---|---|
| Read 1 | AGATCGGAAGAGCACACGTCTGAACTCCAGTCA |
| Read 2 | AGATCGGAAGAGCGTCGTGTAGGGAAAGAGTGT |

### Table 3: Changes to the lib/PARTIES/Map.pm file

<table>
<tr><td>old line</td><td>system("$bowtie2 --threads $self->{THREADS}  --quiet --local -x $self->{BT2_INDEX} -1 $self->{FASTQ1} -2 $self->{FASTQ2} -X $self->{MAX_INSERT_SIZE} | $samtools view -uS - | $samtools sort - -o $out_bam > /dev/null 2>&1");</td></tr>
<tr><td>new line</td><td>system("$bowtie2 --threads $self->{THREADS}  --quiet --local -x $self->{BT2_INDEX} -1 $self->{FASTQ1} -2 $self->{FASTQ2} -X $self->{MAX_INSERT_SIZE} | $samtools view - -uS | $samtools sort - -o notneeded.bam > $out_bam ");</td></tr>
</table>

### Table 4: Input Sequences for AlphaFold2 modeling

| | |
|---|---|
| ISWI1 full length | MSNQSDDENEVLQVELASDEEQRAEEEDERIKKLEQDKKSFMSQIKSTGRMNTNIKFDNIESKINTLLENAEKYAMFL<br>LHRHKRTQESKQKVQGQQRGKHRQIVEDGSEEEDFDDTPTVLEKQPTILKGGQLKSYQLTGLNWMISLFEEGINGIL<br>ADEMGLGKTIQTIGFLAFLKEYKKISGPYLIVAPKSTLGNWMREFKIWMPCMRVVKLIAIKEERDEILNRYFQPGKFDV<br>CLTSYEGVNICLKHIRRFQYKYIIIDEAHKIKNEDAIISQNLRKIRTNYKLLLTGTPLQNTPHELWSLLNYLLPDLFDSSEV<br>FDKWFEVNTEAKLKEGNETIHQDELEQRNLEMVQKFQKILRPFMLRRTKAEVERMLPPKQEIHLFIKMSNLQKSMYQ<br>NILIHNNPHEGDDKGFYMNKLMQLRKICLHPYLFPEVEDKSLPALGEHLVDVSGKMRVLDKFLQKLSEGQHQILIFSQ<br>FTMMLNILEDYCNFRGYEYCRIDGETEIQSRDDQIAEFTAPDSKKFIFLLSTRAGGLGINLATADTVIIYDSDFNPQMDM<br>QAMDRAHRIGQKSRVMVYRMACEHTVEEKIIERQQIKLRWDSLMVQQGRLQQKQNGKLLSKEDLKELTTYGASQIF<br>KLDGDDIKDEDIDILLKRGEQLTKEMNERIEKKFENFKDKVQSLDLGLGQINIFDYFDEAKRNKEDEDALEDALVNHLM<br>QDNKTRNRDKRAMMIGTNSKKIQGKQIKLSEHHLYENKDRLQYLLQKEEDFLAQQKTQKKANENDENVDFGGLTQD<br>ERQEQKRLLETGFKNWNKQEFQDFITANEKYGKDAYEKIQEVIKTKSQDEVKAYAQAFWERIDGLSEKDKIVKQIERG<br>QKLIEQKTNGQKLIEEKCKHFHQPKYELVFTPQLYNKFKSKYFSLENDKFLIYMTNEVGYGNWAQLKQSIRKEPMFRF<br>DHAFKCKSENELKNRVISLVKVLDKEKENNSMGRSLVKNTYIEKPKVLQESQKKKAKNDEEDVQDGSESVKKVKV |
| ISWI1 N-terminus | MSNQSDDENEVLQVELASDEEQRAEEEDERIKKLEQDKKSFMSQIKSTGRMNTNIKFDNIESKINTLLENAEKYAMFL<br>LHRHKRTQESKQKVQGQQRGKHRQIVEDGSEEEDFDDTPTVLEKQPTILKGGQLKSYQLTGLNWMISLFEEGINGIL<br>ADEMGLGKTIQTIGFLAFLKEYKKISGPYLIVAPKSTLGNWMREFKIWMPCMRVVKLIAIKEERDEILNRYFQPGKFDV<br>CLTSYEGVNICLKHIRRFQYKYIIIDEAHKIKNEDAIISQNLRKIRTNYKLLLTGTPLQNTPHELWSLLNYLLPDLFDSSEV<br>FDKWFEVNTEAKLKEGNETIHQDELEQRNLEMVQKFQKILRPFMLRRTKAEVERMLPPKQEIHLFIKMSNLQKSMYQ<br>NILIHNNPHEGDDKGFYMNKLMQLRKICLHPYLFPEVEDKSLPALGEHLVDVSGKMRVLDKFLQKLSEGQHQILIFSQ<br>FTMMLNILEDYCNFRGYEYCRIDGETEIQSRDDQIAEFTAPDSKKFIFLLSTRAGGLGINLATADTVIIYDSDFNPQMDM<br>QAMDRAHRIGQKSRVMVYRMACEHTVEEKIIERQQIKLRWDSLMVQQGRLQ |
| ISWI1 C-terminus | QKQNGKLLSKEDLKELTTYGASQIFKLDGDDIKDEDIDILLKRGEQLTKEMNERIEKKFENFKDKVQSLDLGLGQINIFD<br>YFDEAKRNKEDEDALEDALVNHLMQDNKTRNRDKRAMMIGTNSKKIQGKQIKLSEHHLYENKDRLQYLLQKEEDFLA<br>QQKTQKKANENDENVDFGGLTQDERQEQKRLLETGFKNWNKQEFQDFITANEKYGKDAYEKIQEVIKTKSQDEVKA<br>YAQAFWERIDGLSEKDKIVKQIERGQKLIEQKTNGQKLIEEKCKHFHQPKYELVFTPQLYNKFKSKYFSLENDKFLIYM<br>TNEVGYGNWAQLKQSIRKEPMFRFDHAFKCKSENELKNRVISLVKVLDKEKENNSMGRSLVKNTYIEKPKVLQESQK<br>KKAKNDEEDVQDGSESVKKVKV |
| ICOP1 | MDNKENEKQAKLKEFQRRFPNYMNGKKVIFPIMDEIIVQFSQIFPQSNQFGKMREGNVIKTFSPIPLNQQIEITTFINAF<br>PYNQQFAAMSDSQNQFQLLLSPLLKLTGHLIKDYHTNATFSNSYQYSTFDTQEQSDRINFKLAAFIVEDIFKYKMGLLK<br>PIDIAKQRQVKDEIKKKGNKPQKMSLLSIVEKQQEEEKDLNQDLTIQSQKYEFENYLDPTCQFFWKIAYESFNKIIKKS<br>QRTKLQDMDIEQDSDEQPETIQNVNKETLSQDNIEMRRQQIISKYQQLCKLKDQSKKKSKQTKQYSQIKSFKIKDRYT<br>DLEMLRFNNLFKFLVQNWPSFLLQSIKLPYVQSLFTDQELRNIKSIGQNELGYFGLKAKQRADITSNLIEGVRETDSFK<br>VIIEYRQEVTEAVGLVLENVQDELNSINQSLQKKDSQYTQQQQQQYRKVYKQYLQLVEFNRLFINGCLYLGSDIHGFD<br>YHIFSNDIDHIYQNNGSEWRVLDENQVQQLFKTLNVCGVKERELQTNIQKLMACELFNDQETKELITIKNVEQSQVQA<br>GNRSPKQLIVKILLEVVQKYTDILMVRKLRWESYKIREKFQNTIKTLENPLDMVDFMKILIEQFETAQVLIIDQQKMQNG<br>SQYDQRDLKEFQQRLRIYENKLKGLKEPEKILFYDTQLFQMMESREHIKPNGVKCNTKFWQQSLGLEVKEALMNFA<br>NKVDKEHQQYDVVFMASTLLLAVQEYELSSSAQDNEDDELLRQIVKDVKVDNSNFPSKIDNHQKNNQIIELD |
| ICOP2 | MDNKENEKQAKLKEFQRRFPNYMNGKKVIFPIMDEIIVQFSQIFPQSNQFGKMREGNVIKTFSPIPLNQQIEITTFINAF<br>PYNQQFAAMSDSQNQFQLLLSPLLKLTGHLIKDYHSNATFSNSYQYSTFDTQEQSDRINFKLTAFIIEDIFKYKMGLLK<br>PIDIAKQRQVKDDIKKKANKAQKISLLTMVEKQQQQEEEKDVNQDLTLQSQKYEFENYLDPTCQSFWKIAYEAFNKIIK<br>KSQRTKLQDMDIEQESDEQPDTIQNVNNQKGNLNQESIELRRQQIISKYQQLGKLKDQNKKKSKSTKQYTQIKSFKIK<br>DKYTDLEMLRFNNLFKFLIQNWPAMLLQSIKLPYVQSLFNDQELRNIKSIGQNDLGYFGLKAKQRADITSNLIEGVRET<br>DSFKIIIEYRQEVTEAVGLVLENVQDELNSINQALQKKDSQYTLQQQQQYRKIYKQYLQLVEFSRLFINGCLYLGSDIH<br>GYDYHIFSNDIDHIYQNNGSEWRVLDENQVQQLFKTLNVCGVKERELQTNIQKLMACELFNDQDTKELITIKNVEQSQ<br>VQAGNRSPKQLIVKILLEVVQKYTDTLMIRKLRWESYKIREKFQNTIKTLENPLDMVDFMKILIEQFETAQVLVIDQQKM<br>QNGNQYDQKDLKEFQQRIRIYENKLQGLKEPEKILFYDSQLFQLMESKEYIKPNGIKSNTKFWQQSLGVEVKEALMN<br>FANKVDKENQQYDVVFMASTLLLAVQEYELSSSSQDNEDDELLRQIVKEVKFDNQNLLLKTDNHQVNNQIIELD |
| Ptiwi01 | MFQNIQLKANFHEMRLNPSRPVYQYKLEITDSSPEKVSEALKKFRPQLQTQLILFMSLNQNIYSPKLIQEADNGLVLGS<br>LSGNETNQDTATLKLVGKIENKADLNIIISRLFKQVIRSQMQMVSVGNKGQKLFWSSRAQQFKDQNLEIWPGVECIFR<br>PGEGGAQNPTLVIDCAFKMLRYRSALEELNQTRNPACIQDQIVMTTYNKKFYKVEAVDVNLKPASTFTNEKGETISFA<br>QYYEQRYKVKVDGNQPLIRATVRSKQDKTEKTIHLIPQLCQLTGLTDAIRNDFNAMKNLAVVTKPGADQRMKMAQEF<br>ANQLASTEIVNKKLGTKRQIFKEWGVEINPGSMDVPARRIHPGNMLMGNGLKLDLSSPQTNLDRQTQTQMFSTPPQ<br>QLILGIIYNKKTGQQTMDSLMQNFQAACNDFKFQAFMAPKVFPIEQDRDEDLERVLDGFQKQAEANKAKVGFLLFLL<br>PGQKKKARLYKTAKKISMQKFGCASQVVVEKTLAKNTRSIVNKILIQLNAKVGGTPWAIDSLPTTFQNQPTMICGTDC<br>FVKSGRKNQLAFCSTVDRNLSRYYSQVVTSGEFSQHLQQVFKASLLAFKEQNGIFPKLIIIYRDGVGDGQQAVVLANE<br>LPQYKQALEELQITDTKISLVVCNKRVSAKFYTGGNARPDNPQPGTCVDNPKVVEQSNPNFYLISQVTRQGTVTPSL<br>YKIIHSDQAGLDDDIKVLTFKLCWLFYNFTGSIKIPAPVRYAHCLCNFIGDNYDDRDQVKFLPLPDLVKQKVLFYI |

### Table 5: Codon Optimized sequences for recombinant protein expression

| | |
|---|---|
| ISWI1 | MSNQSDDENEVLQVELASDEEQRAEEEDERIKKLEQDKKSFMSQIKSTGRMNT<br>NIKFDNIESKINTLLENAEKYAMFLLHRHKRTQESKQKVQGQQRGKHRQIVEDGS<br>EEEDFDDTPTVLEKQPTILKGGQLKSYQLTGLNWMISLFEEGINGILADEMGLGK<br>TIQTIGFLAFLKEYKKISGPYLIVAPKSTLGNWMREFKIWMPCMRVVKLIAIKEERD<br>EILNRYFQPGKFDVCLTSYEGVNICLKHIRRFQYKYIIIDEAHKIKNEDAIISQNLRKI<br>RTNYKLLLTGTPLQNTPHELWSLLNYLLPDLFDSSEVFDKWFEVNTEAKLKEGNE<br>TIHQDELEQRNLEMVQKFQKILRPFMLRRTKAEVERMLPPKQEIHLFIKMSNLQK<br>SMYQNILIHNNPHEGDDKGFYMNKLMQLRKICLHPYLFPEVEDKSLPALGEHLVD<br>VSGKMRVLDKFLQKLSEGQHQILIFSQFTMMLNILEDYCNFRGYEYCRIDGETEI<br>QSRDDQIAEFTAPDSKKFIFLLSTRAGGLGINLATADTVIIYDSDFNPQMDMQAMD<br>RAHRIGQKSRVMVYRMACEHTVEEKIIERQQIKLRWDSLMVQQGRLQQKQNGK<br>LLSKEDLKELTTYGASQIFKLDGDDIKDEDIDILLKRGEQLTKEMNERIEKKFENFK<br>DKVQSLDLGLGQINIFDYFDEAKRNKEDEDALEDALVNHLMQDNKTRNRDKRAM<br>MIGTNSKKIQGKQIKLSEHHLYENKDRLQYLLQKEEDFLAQQKTQKKANENDENV<br>DFGGLTQDERQEQKRLLETGFKNWNKQEFQDFITANEKYGKDAYEKIQEVIKTK<br>SQDEVKAYAQAFWERIDGLSEKDKIVKQIERGQKLIEQKTNGQKLIEEKCKHFHQ<br>PKYELVFTPQLYNKFKSKYFSLENDKFLIYMTNEVGYGNWAQLKQSIRKEPMFRF<br>DHAFKCKSENELKNRVISLVKVLDKEKENNSMGRSLVKNTYIEKPKVLQESQKKK<br>AKNDEEDVQDGSESVKKVKV |
| ICOP1 | MDNKENEKQAKLKEFQRRFPNYMNGKKVIFPIMDEIIVQFSQIFPQSNQFGKMRE<br>GNVIKTFSPIPLNQQIEITTFINAFPYNQQFAAMSDSQNQFQLLLSPLLKLTGHLIK<br>DYHTNATFSNSYQYSTFDTQEQSDRINFKLAAFIVEDIFKYKMGLLKPIDIAKQRQ<br>VKDEIKKKGNKPQKMSLLSIVEKQQEEEKDLNQDLTIQSQKYEFENYLDPTCQFF<br>WKIAYESFNKIIKKSQRTKLQDMDIEQDSDEQPETIQNVNKETLSQDNIEMRRQQI<br>ISKYQQLCKLKDQSKKKSKQTKQYSQIKSFKIKDRYTDLEMLRFNNLFKFLVQNW<br>PSFLLQSIKLPYVQSLFTDQELRNIKSIGQNELGYFGLKAKQRADITSNLIEGVRET<br>DSFKVIIEYRQEVTEAVGLVLENVQDELNSINQSLQKKDSQYTQQQQQQYRKVY<br>KQYLQLVEFNRLFINGCLYLGSDIHGFDYHIFSNDIDHIYQNNGSEWRVLDENQV<br>QQLFKTLNVCGVKERELQTNIQKLMACELFNDQETKELITIKNVEQSQVQAGNRS<br>PKQLIVKILLEVVQKYTDILMVRKLRWESYKIREKFQNTIKTLENPLDMVDFMKILIE<br>QFETAQVLIIDQQKMQNGSQYDQRDLKEFQQRLRIYENKLKGLKEPEKILFYDTQ<br>LFQMMESREHIKPNGVKCNTKFWQQSLGLEVKEALMNFANKVDKEHQQYDVVF<br>MASTLLLAVQEYELSSSAQDNEDDELLRQIVKDVKVDNSNFPSKIDNHQKNNQII<br>ELD |
| ICOP2 | MDNKENEKQAKLKEFQRRFPNYMNGKKVIFPIMDEIIVQFSQIFPQSNQFGKMRE<br>GNVIKTFSPIPLNQQIEITTFINAFPYNQQFAAMSDSQNQFQLLLSPLLKLTGHLIK<br>DYHSNATFSNSYQYSTFDTQEQSDRINFKLTAFIIEDIFKYKMGLLKPIDIAKQRQV<br>KDDIKKKANKAQKISLLTMVEKQQQQEEEKDVNQDLTLQSQKYEFENYLDPTCQ<br>SFWKIAYEAFNKIIKKSQRTKLQDMDIEQESDEQPDTIQNVNNQKGNLNQESIELR<br>RQQIISKYQQLGKLKDQNKKKSKSTKQYTQIKSFKIKDKYTDLEMLRFNNLFKFLI<br>QNWPAMLLQSIKLPYVQSLFNDQELRNIKSIGQNDLGYFGLKAKQRADITSNLIEG<br>VRETDSFKIIIEYRQEVTEAVGLVLENVQDELNSINQALQKKDSQYTLQQQQQYR<br>KIYKQYLQLVEFSRLFINGCLYLGSDIHGYDYHIFSNDIDHIYQNNGSEWRVLDEN<br>QVQQLFKTLNVCGVKERELQTNIQKLMACELFNDQDTKELITIKNVEQSQVQAGN<br>RSPKQLIVKILLEVVQKYTDTLMIRKLRWESYKIREKFQNTIKTLENPLDMVDFMKI<br>LIEQFETAQVLVIDQQKMQNGNQYDQKDLKEFQQRIRIYENKLQGLKEPEKILFY<br>DSQLFQLMESKEYIKPNGIKSNTKFWQQSLGVEVKEALMNFANKVDKENQQYD<br>VVFMASTLLLAVQEYELSSSSQDNEDDELLRQIVKEVKFDNQNLLLKTDNHQVN<br>NQIIELD |

#### Table 6: Samples used for nucleosome density analyses

| Sample | Reference |
|---|---|
| *ICOP1/2/PGM*-KD | This study |
| *ND7/PGM*-KD (control for this study) | This study |
| *ISWI1/PGM*-KD | (Singh et al. 2022) |
| *NOWA1/2/PGM*-KD | (Singh et al. 2022) |
| *PTCAF1/PGM*-KD | (Wang et al. 2022) |
| *empty vector/PGM*-KD<br>(control for *PTCAF1/PGM*-KD & *NOWA1/2/PGM*-KD) | (Singh et al. 2022; Wang et al. 2022) |

RNA-guided PRC2 complex eliminates DNA as an extreme form of transposon silencing. *Cell Rep* **40**: 111263.
